# Pathformer: a biological pathway informed Transformer integrating multi-omics data for disease diagnosis and prognosis

**DOI:** 10.1101/2023.05.23.541554

**Authors:** Xiaofan Liu, Yuhuan Tao, Zilin Cai, Pengfei Bao, Hongli Ma, Kexing Li, Mengtao Li, Yunping Zhu, Zhi John Lu

**Author notes:** To whom correspondence should be addressed: Zhi John Lu; Mengtao Li; Yunping Zhu.

## Abstract

Multi-omics data provide a comprehensive view of gene regulation at multiple levels, which is helpful in achieving accurate diagnosis of complex diseases like cancer. To integrate various multi-omics data of tissue and liquid biopsies for disease diagnosis and prognosis, we developed a biological pathway informed Transformer, Pathformer. It embeds multi-omics input with a compacted multi-modal vector and a pathway-based sparse neural network. Pathformer also leverages criss-cross attention mechanism to capture the crosstalk between different pathways and modalities. We first benchmarked Pathformer with 18 comparable methods on multiple cancer datasets, where Pathformer outperformed all the other methods, with an average improvement of 6.3%-14.7% in F1 score for cancer survival prediction and 5.1%-12% for cancer stage prediction. Subsequently, for cancer prognosis prediction based on tissue multi-omics data, we used a case study to demonstrate the biological interpretability of Pathformer by identifying key pathways and their biological crosstalk. Then, for cancer early diagnosis based on liquid biopsy data, we used plasma and platelet datasets to demonstrate Pathformer’s potential of clinical applications in cancer screen. Moreover, we revealed deregulation of interesting pathways (e.g., scavenger receptor pathway) and their crosstalk in cancer patients’ blood, providing new candidate targets for cancer microenvironment study.

## Introduction

Comparing to single type of data, multi-omics data provide a more comprehensive view of gene regulation^1^. Therefore, integrating multi-omics data from tissue and liquid biopsies would be helpful in addressing challenges in disease diagnosis and prognosis, such as deregulated network between different types of molecules and data noise caused by patients’ heterogeneity^2^. To integrate multi-omics data of cancer, several supervised methods have been developed, such as mixOmics^3^, liNN^4^, eiNN^5^, liCNN^6^, eiCNN^7^, MOGONet^8^ and MOGAT^9^. Later, the performance and interpretability of multi-omics data integration were further improved using deep learning models informed by biological pathways. For instance, a pathway-associated sparse deep neural network (PASNet) was utilized to predict the prognosis of glioblastoma multiforme (GBM) patients^10^. Recently, P-NET, a sparse neural network integrating multiple molecular features based on a multilevel view of biological pathways, was introduced to predict subtype and survival of prostate cancer patients^11^. In addition, PathCNN based on a convolutional neural network (CNN) was developed to predict the prognosis of GBM patients using principal component analysis (PCA) to define image-like multi-omics pathways^12^.

These pathway-informed deep learning methods did not consider the crosstalk between omics and between pathways, although the crosstalk holds biological significance as well as pathway itself^13-15^. Meanwhile, the criss-cross attention mechanism of the Transformer would be very useful to capture the crosstalk information^16^. However, incorporating multi-omics data and their crosstalk information in a Transformer is very challenging: when processing multi-omics data, the multi-modal features are usually multiplied by the number of genes (tens of thousands), producing an extremely long input that is not acceptable by a common Transformer model (usually less than 512). Meanwhile, certain embedding methods for biological data, such as discretization and linear transformation, were introduced in the previous Transformer models^17-19^, while biological information was largely lost during these kinds of embedding.

In order to integrate multi-omics data by embedding biological pathway crosstalk without information loss, we introduce a Transformer model, Pathformer, with three key steps to address the above problems. First, it ingeniously concatenates different modalities into a novel compacted multi-modal vector for each gene, which not only saves valuable information but also shortens the input. Second, Pathformer utilizes a sparse neural network based on prior pathway knowledge to transform gene embeddings into pathway embeddings. Third, Pathformer naturally incorporates pathway crosstalk network into a Transformer model with bias to enhance the exchange of information between different pathways and between different modalities (e.g., omics) as well.

Here, we first benchmarked Pathformer and 18 other integration methods in various classification tasks, using multiple cancer tissue datasets from TCGA. Then, we used Pathformer to integrate various multi-omics data from tissue and liquid biopsies. Through case studies on survival prediction of breast cancer and noninvasive diagnosis of pan-cancer, we revealed interesting pathways, genes, and regulatory mechanisms related to cancer in human tissue and plasma, demonstrating the prediction accuracy and biological interpretability of Pathformer in various clinical applications.

## Materials and methods

### Overview of Pathformer

Pathformer is mainly designed to integrate various multi-omics data from tissue and liquid biopsies, which can be used for different classification tasks in disease diagnosis and prognosis, such as cancer early detection, cancer staging and survival prediction (**Fig. 1a**). It has 6 modules: 1) biological pathway and crosstalk network calculation module, 2) multi-omics data input module, 3) biological multi-modal embedding module (key module), 4) Transformer module with pathway crosstalk network bias, 5) classification module and 6) biological interpretability module.

**Figure 1.**
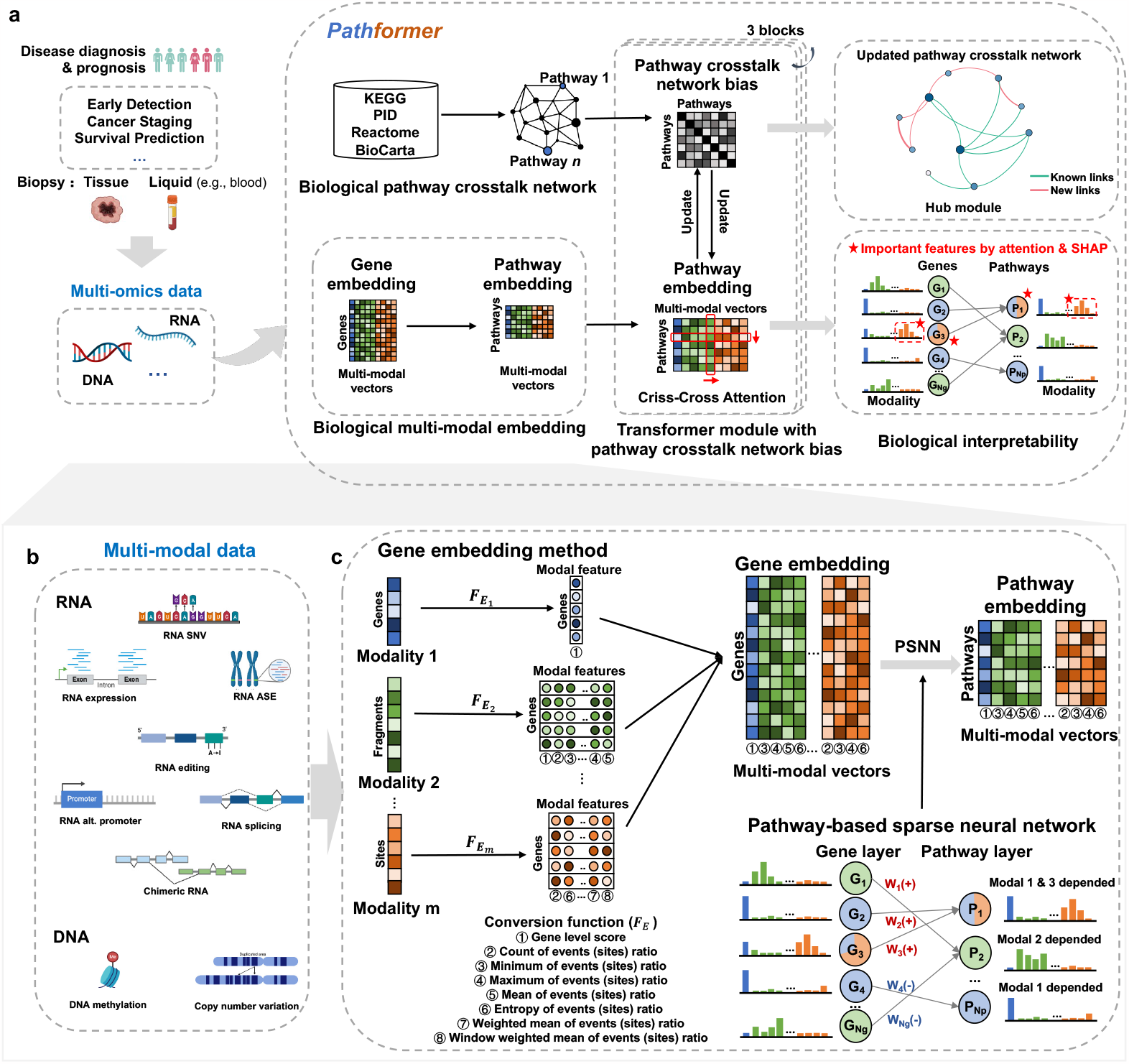
Overview of Pathformer. Schematic of Pathformer (**a**), which integrates multi-oimcs data of tissue and liquid biopsies for disease diagnosis and prognosis. Pathformer has 6 modules: 1) biological pathway and crosstalk network calculation module, 2) multi-omics data input module (**b**), 3) biological multi-modal embedding module (**c**), 4) Transformer module with pathway crosstalk network bias, 5) classification module and 6) biological interpretability module. *F*_*E*_, conversion function in the gene embedding; G, gene; P, pathway; W, weight of pathway-based sparse neural network.

Pathformer combines prior biological pathway information (module 1, **Fig. 1a**) with multi-modal data (module 2, **Fig. 1b**) for disease diagnosis and prognosis. It introduces a new embedding method to incorporate biological multi-modal data at both gene level and pathway level: it initiates the process by uniformly transforming different modalities to the gene level through a series of statistical indicators, then concatenates these modalities into compacted multi-modal vectors to define gene embedding, and employs a sparse neural network based on the gene-to-pathway mapping to transform gene embedding into pathway embedding (module 3, **Fig. 1c**). Pathformer then enhances the fusion of information between various modalities and pathways by combining pathway crosstalk networks with Transformer encoder (module 4, **Fig. 1a, Supplementary Fig. 1**). Finally, a fully connected layer serves as the classifier for different downstream classification tasks (module 5). In addition, Pathformer uses a biological interpretable module with attention weights and SHapley Additive exPlanations^20^ values to identify important genes, pathways, modalities, and their crosstalk or regulation (module 6).

These 6 modules are described in detail below.

### Module 1: curation of biological pathways and calculation of initial crosstalk network

We curated 2,289 biological pathways from four public databases including Kyoto Encyclopedia of Genes and Genomes database (KEGG)^21^, Pathway Interaction database (PID)^22^, Reactome database (Reactome)^23^, and BioCarta Pathways database (BioCarta)^24^. Then, we filtered these pathways by three criteria: the gene number, the overlap ratio with other pathways (the proportion of genes in the pathway that are also present in other pathways), and the number of pathway subsets (the number of pathways included in the pathway). Following the principle of moderate size and minimal overlap with other pathway information, we selected 1,497 pathways with gene number between 15 and 100, or gene number greater than 15 and overlap ratio less than 1, or gene number greater than 15 and the number of pathway subsets less than 5. Next, we used *BinoX*^25^ to calculate the crosstalk relationship of pathways and build a pathway crosstalk network with adjacency matrix 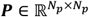, *N*_*p*_=1.497 (more details in **Supplementary Note 1**).

### Modules 2 and 3: multi-omics data input and multi-modal embedding

Biological multi-modal data preprocessing and embedding method are two key modules of Pathformer (**Fig. 1b, c**). In module 2 (**Fig. 1b**), to capture more comprehensive regulatory information, we expand biological multi-omics data into multi-modal data, including not only data from different omics sources but also variant features of the same omics, such as RNA splicing, RNA editing, RNA alternative promoter, and so on. To obtain multi-modal data, we use standardized bioinformatics pipeline to calculate different omics or variant features of the same omics from raw sequence reads (more details in **Supplementary Note 5**). These multi-modal data have different dimensions, including nucleotide level, fragment level, and gene level. For example, Pathformer’s input for cancer tissue datasets from Cancer Genome Atlas (TCGA)^26^ includes gene-level RNA expression, fragment-level DNA methylation, and both fragment-level and gene-level DNA CNV. Modalities and their dimensions level for different datasets are described in **Supplementary Table 1**.

In module 3 (**Fig. 1c**), we proposed a new biological multi-modal embedding method of Pathformer, which consists of gene embedding ***E***_*G*_ and pathway embedding ***E***_*P*_. We represented biological multi-modal input matrix of a sample as ***M***, described as follows:

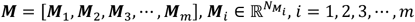

where *m* is the number of modalities, and 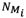 is the length of input for modality *i*, likes the number of genes for RNA expression, the number of editing sites for RNA editing, and the number of CpG islands for DNA methylation. Next, we first use a series of statistical indicators to convert different modalities into gene level modal features, and then concatenate these modal features into a compressed multi-modal vector as gene embedding ***E***_*G*_, which are calculated as follows:

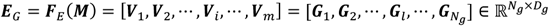

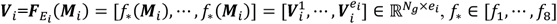

where 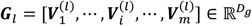 is gene embedding of the *l*th gene and is a compacted multi-modal vector; ***V***_*i*_ is the modal features matrix of modality *i*; 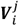 is the *j*th dimension of modal features matrix for modality *i*; *N*_*g*_ is the number of genes, *D*_*g*_ = *e*_1_ + *e*_2_ + … + *e*_*m*_ is the dimension of gene embedding; *e*_*_ is the dimension of modal features matrix for modality *i*; ***F***_*E*_ is the conversion function, which uses statistical indicators to uniformly convert different modalities into gene level; 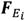 is the conversion function of modality *i*, and each modality’s function is constructed from distinct statistical indicator functions *f*_*_ (more details in **Supplementary Table 1**). These statistical indicator functions include gene level score (*f*_1_), count (*f*_2_), minimum (*f*_3_), maximum (*f*_4_), mean (*f*_5_), entropy (*f*_6_), weighted mean in whole gene (*f*_7_) and weighted mean in window (*f*_8_), formulas of which are in **Supplementary Note 2**.

Subsequently, we used the known gene-pathway mapping relationship to develop a sparse neural network based on prior pathway knowledge (PSNN) to transform gene embedding ***E***_*G*_ into pathway embedding ***E***_*P*_, as described below:

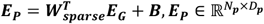

where *N*_*p*_ is the number of pathways, *D*_*p*_ = *D*_*g*_ is the dimension of pathway embedding, 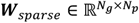 is a learnable sparse weight matrix, and ***B*** is a bias term. ***W***_*sparse*_ is constructed based on the known relationship between pathways and genes. When the given gene and the pathway are irrelevant, the corresponding element of ***W***_*sparse*_ will always be 0. Otherwise, it needs to be learned through training. Therefore, pathway embedding is a dynamic embedding method. The PSNN can not only restore the mapping relationship between genes and pathways, but also capture the different roles of different genes in pathways, and can preserve the complementarity of different modalities. Additionally, this biological multi-modal embedding step does not require additional gene selection, thereby avoiding bias and overfitting problems resulting from artificial feature selection.

### Module 4: Transformer module with pathway crosstalk network bias

We developed the Transformer module based on criss-cross attention (CC-attention) with bias for data fusion of pathways, modalities, and their crosstalk (**Supplementary Fig. 1**). This module has 3 blocks, each containing multi-head column-wise self-attention (col-attention), multi-head row-wise self-attention (row-attention), layer normalization, GELU activation, residual connection, and network update. Particularly, col-attention is used to enhance the exchange of information between pathways, with the pathway crosstalk network matrix serving as the bias for col-attention to guide the flow of information. Row-attention is employed to facilitate information exchange between different modalities, and the updated pathway embedding matrix is used to update the pathway crosstalk network matrix by calculating the correlation between pathways.

Multi-head column-wise self-attention contains 8 heads, each head is a mapping of ***Q***_**1**_,***K***_**1**_,***V***_**1**_,***P***, which are query vector, key vector, and value vector of pathway embedding ***E***_*P*_ and pathway crosstalk network matrix ***P***, respectively. First, we represented the *h*th column-wise self-attention by 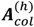, calculated as follows:

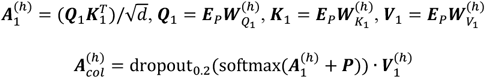

where *h* = 1,2, …, *H* is the *h*th head; *H* is the number of heads; 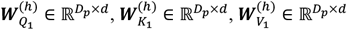 are the weight matrices as parameters; *d* is the attention dimension; dropout_0.2_ is a dropout neural network layer with a probability of 0.2; and softmax is the normalized exponential function.

Next, we merged multi-head column-wise self-attention and performed a series of operations as follows:

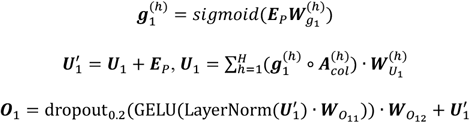

where *h* = 1,2, …, *H* is the *h*th head; *H* is the number of heads; ∘ is the matrix dot product; 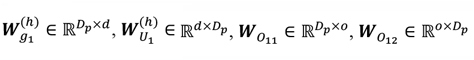 are the weight matrices as parameters; *o* is a constant; LayerNorm is the layer normalization function; GELU is the distortion of RELU activation function; and dropout_0.2_is a dropout neural network layer with a probability of 0.2.

Multi-head row-wise self-attention enables information exchange between different modalities. It is a regular dot-product attention. It also contains 8 heads, and the *h*th row-wise self-attention, i.e., 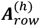, is calculated as follows:

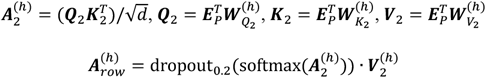

where *h* = 1,2, …, h is the *h*th head; *H* is the number of heads; 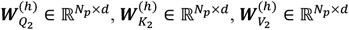 are the weight matrices as parameters; *d* is the attention dimension; dropout_0.2_ is a dropout neural network layer with a probability of 0.2; and softmax is the normalized exponential function.

Subsequently, we merged multi-head row-wise self-attention and performed a series of operations. The formulas are as follows:

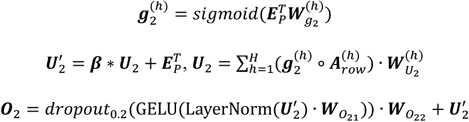

where *h* = 1,2, …, h is the *h*th head; *H* is the number of heads; ∘ is the matrix dot product; 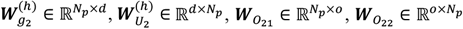 are the weight matrices as parameters; *o* is a constant; ***β*** is a constant coefficient for row-attention; LayerNorm is the layer normalization function; GELU is the distortion of RELU activation function; and *dropout*_0.2_ is a dropout neural network layer with a probability of 0.2. ***O***_2_ is pathway embedding input of the next Transformer block. In other words, when ***E***_*P*_ is 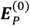, ***O***_2_ is 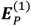, superscripts with parenthesis represent data at different block.

Then, we used the updated pathway embedding ***O***_2_ to update the pathway crosstalk network. We exploited the correlation between embedding vectors of two pathways to update the corresponding element of the pathway crosstalk network matrix. The formula is as follows:

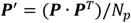

where ***P***′ is the updated pathway crosstalk network matrix of next Transformer block. In other words, when ***P***′ is ***P***^(1)^, ***P*** is ***P***^(0)^, superscripts with parenthesis represent data at different block.

### Module 5: classification module

Given the classification tasks in disease diagnosis and prognosis, we used the fully connected neural network as the classification module to transform pathway embedding encoded by the Transformer module into the probability for each label. Three fully connected neural networks each have 300, 200, and 100 neurons, with dropout probability *dropout*_*c*_, which is hyperparameter. More details of the classification module are described in **Supplementary Note 3**.

### Model 6: biological interpretability module

The biological interpretable module enables us to calculate the contribution of each modality, identify important pathways and their key genes, and uncover the most critical pathway crosstalk subnetworks.

To calculate the contribution of each omics and each modality, we first integrated all matrices of row-attention maps into one matrix by element-wise average. Then, we averaged this average row-attention matrix along with columns as the attention weights of modalities. More details are described in **Supplementary Note 4**.

To identify important pathways and their key genes, we used SHapley Additive exPlanations^20^ (SHAP value) to calculate the contribution of each feature. It is an additive explanation model inspired by coalitional game theory, which regards all features as “contributors”. SHAP value is the value assigned to each feature, which explains the relationship between modalities, pathways, genes and classification, implemented by “SHAP” package of *Python* v3.6.9. Then, pathways with the top 15 SHAP values in the classification task are considered as important pathways. For each pathway, genes with top 5 SHAP values are considered as the key genes of the pathway. The core modality on which one gene depends indicates that the SHAP value of that gene ranks higher on this modality than on the others. More details are described in **Supplementary Note 4**.

Particularly, the pathway crosstalk network matrix is used to guide the direction of information flow, and updated according to updated pathway embedding in each Transformer block. Therefore, the updated pathway crosstalk network contains not only the prior information in the initial network (module 1) but also the multi-modal data information derived from the Transformer module (module 4), which represents the specific regulatory mechanism in each classification task. We defined the sub-network score through SHAP value of each pathway in sub-network, so as to find foremost sub-network for prediction, that is, hub module of the updated pathway crosstalk network. The calculation of the sub-network score can be divided into four steps: average pathway crosstalk network matrix calculation, network pruning, sub-network boundary determination, and score calculation. More details of sub-network score calculations are described in **Supplementary Note 4**.

### Experimental settings

#### Data collection and preprocessing

We assayed both tissue biopsy and liquid biopsy data in this study. First, for benchmark testing on cancer diagnosis and prognosis, we collected multiple datasets of different cancer types from TCGA (tissue data) to evaluate the classification performance, including 10 datasets for early- and late-stage classification, and 10 datasets for low- and high-risk survival classification (**Supplementary Fig. 2**). In addition, we also collected and processed two types of body fluid datasets: the plasma dataset (373 samples assayed by total cell-free RNA-seq^27,28^) and the platelet dataset (918 samples assayed by blood platelet RNA-seq^29,30^). Through our biological information pipeline, 3 and 7 biological modalities were derived from the TCGA (tissue biopsy) datasets and the liquid biopsy datasets, respectively. More details are described in **Supplementary Notes 5 and 7**.

#### Model training and test

We implemented Pathformer’s network architecture using the “PyTorch” package in *Python* v3.6.9, and our codes can be found in the GitHub repository (https://github.com/lulab/Pathformer). For model training and test, we used 5-fold cross-validation, and repeated it twice by shuffling. Before evaluating the performance on the test set, we optimized hyperparameters (e.g., learning rate, dropout probability of classification and constant coefficient for row-attention) and epoch numbers inside the training set only. More details of model training and test are described in **Supplementary Note 6**.

#### Evaluation criteria

When evaluating the classification performance, we used at least three evaluation indicators, area under the receiver operating characteristic curve (AUC), weighted-averaged F1 score (F1score_weighted), and macro-averaged F1 score (F1score_macro). Notably, we prioritized F1score_macro as the main evaluation criterion in this paper. This choice stems from the imbalance of sub-classes in our data, where F1score_macro stands out as a fairer and more robust indicator compared to other metrics such as AUC.

## Results

### Benchmark of Pathformer and 18 multi-omics data integration methods using TCGA data

We conducted a meticulous benchmark of Pathformer and other 18 multi-omics data integration methods for various classification tasks in cancer diagnosis and prognosis (**Fig. 2**). These methods can be categorized into 3 types. Type I includes early and late integration methods based on conventional classifiers, such as support vector machine (SVM), logistic regression (LR), random forest (RF), and extreme gradient boosting (XGBoost). Type II includes partial least squares-discriminant analysis (PLSDA) and sparse partial least squares-discriminant analysis (sPLSDA) of mixOmics^3^. Type III consists of deep learning-based integration methods, i.e., eiNN^5^, liNN^4^, eiCNN^7^, liCNN^6^, MOGONet^8^, MOGAT^9^, P-NET^11^ and PathCNN^12^. Among these, eiNN and eiCNN are early integration methods based on NN and CNN; liNN and liCNN are late integration methods based on fully connected neural network (NN) and convolutional neural network (CNN); MOGONet and MOGAT are multi-modal integration methods based on graph neural network; P-NET and PathCNN are representative multi-modal integration methods that combines pathway information.

**Figure 2.**
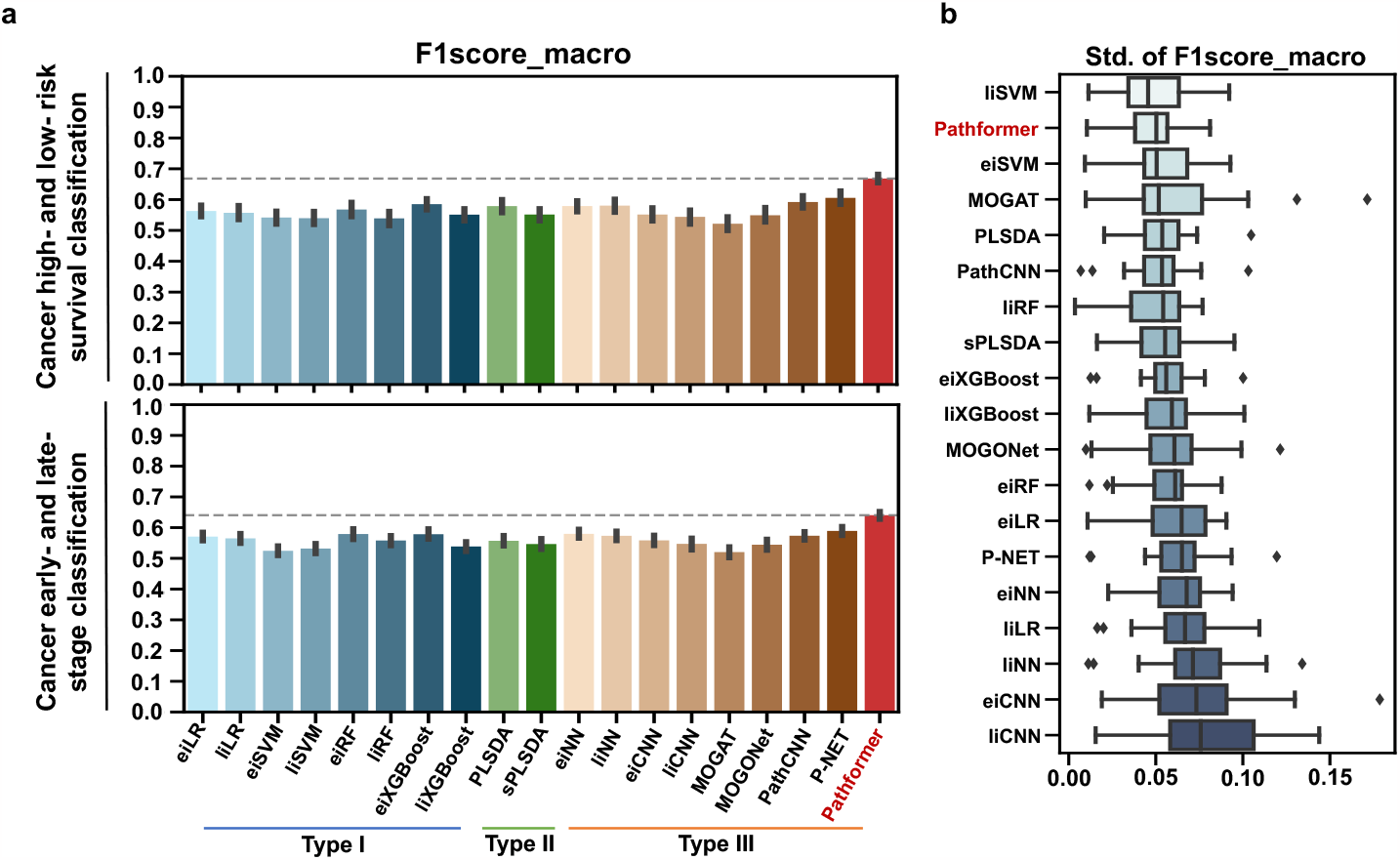
Performance comparison among multiple multi-omics data integration methods. Average macro-averaged F1 score (**a**) and its standard deviation (**b**) are shown for each method on the TCGA datasets (all cancer types) for cancer low- and high-risk survival classification and early- and late-stage classification, respectively. Error bars are from 5-fold cross-validation repeated twice (10 values) of all datasets.

To evaluate the performance, we tested the methods on multiple TCGA datasets for two tasks, cancer survival prediction and cancer staging. DNA methylation, DNA copy number variation (CNV), and RNA expression were used as input. The results of optimal hyperparameter combination for each dataset are listed in **Supplementary Table 2**. Considering the numbers of sub-classes in the TCGA data are imbalanced, we utilized the macro-averaged F1 score as the primary evaluation metric for hyperparameter optimization and performance evaluation (**Fig. 2 and Tables 1-2**). We also list the other evaluation indicators (e.g., AUC) in **Supplementary Table 3**.

**Table 1.** Performance comparison among multiple multi-omics data integration methods for cancer survival prediction.

|  | BRCA<sup>1</sup> | KIRC | LUAD | LUSC | HNSC | BLCA | LIHC | SKCM | LGG | Pan-cancer<sup>2</sup> |
| --- | --- | --- | --- | --- | --- | --- | --- | --- | --- | --- |
| eiLR | 0.571 | 0.584 | 0.530 | 0.520 | 0.529 | 0.479 | 0.497 | 0.557 | 0.689 | 0.674 |
| eiRF | 0.494 | 0.636 | 0.443 | 0.477 | 0.497 | 0.475 | 0.579 | 0.609 | 0.759 | 0.709 |
| eiSVM | 0.467 | 0.656 | 0.487 | 0.471 | 0.466 | 0.459 | 0.436 | 0.630 | 0.640 | 0.705 |
| eiXGBoost | 0.564 | 0.653 | 0.530 | 0.519 | 0.476 | 0.549 | 0.550 | 0.594 | 0.723 | 0.694 |
| liLR | 0.569 | 0.619 | 0.479 | 0.481 | 0.512 | 0.456 | 0.487 | 0.554 | 0.718 | 0.696 |
| liRF | 0.473 | 0.622 | 0.453 | 0.452 | 0.478 | 0.438 | 0.428 | 0.609 | 0.724 | 0.712 |
| liSVM | 0.468 | 0.614 | 0.438 | 0.508 | 0.470 | 0.475 | 0.439 | 0.561 | 0.714 | 0.710 |
| liXGBoost | 0.523 | 0.585 | 0.481 | 0.491 | 0.483 | 0.504 | 0.500 | 0.532 | 0.715 | 0.697 |
| PLSDA | 0.561 | 0.660 | 0.444 | 0.496 | 0.527 | 0.506 | 0.514 | 0.635 | 0.778 | 0.669 |
| sPLSDA | 0.549 | 0.617 | 0.450 | 0.495 | 0.521 | 0.465 | 0.442 | 0.593 | 0.726 | 0.658 |
| eiNN | 0.573 | 0.640 | 0.503 | 0.483 | 0.537 | 0.497 | 0.593 | 0.583 | 0.695 | 0.687 |
| eiCNN | 0.510 | 0.548 | 0.462 | 0.529 | 0.512 | 0.474 | 0.578 | 0.582 | 0.631 | 0.688 |
| liNN | 0.576 | 0.631 | 0.521 | 0.480 | 0.523 | 0.518 | 0.551 | 0.590 | 0.702 | 0.713 |
| liCNN | 0.536 | 0.596 | 0.504 | 0.501 | 0.487 | 0.460 | 0.465 | 0.489 | 0.706 | 0.697 |
| MOGONet | 0.510 | 0.632 | 0.455 | 0.497 | 0.469 | 0.470 | 0.476 | 0.599 | 0.706 | 0.676 |
| MOGAT | 0.466 | 0.637 | 0.438 | 0.434 | 0.462 | 0.443 | 0.448 | 0.605 | 0.595 | 0.689 |
| PathCNN | 0.558 | 0.643 | 0.522 | 0.560 | 0.525 | 0.486 | 0.598 | 0.551 | 0.766 | 0.712 |
| P-NET | 0.640 | 0.661 | 0.551 | 0.509 | 0.559 | 0.470 | 0.578 | 0.641 | 0.715 | 0.733 |
| **Pathformer** | **0.673** | **0.688** | **0.633** | **0.615** | **0.606** | **0.601** | **0.651** | **0.694** | **0.787** | **0.735** |

<sup>1</sup> abbreviation of cancer type according to the TCGA terms; <sup>2</sup>pan-cancer dataset contains 33 cancer types of TCGA terms.
10 TCGA datasets are tested. Average macro-averaged F1 scores are listed for the two unbalanced classes, high- and low-risk survival cancer patients. Each value is the mean of 5-fold cross-validation repeated twice (10 values).

**Table 2.**
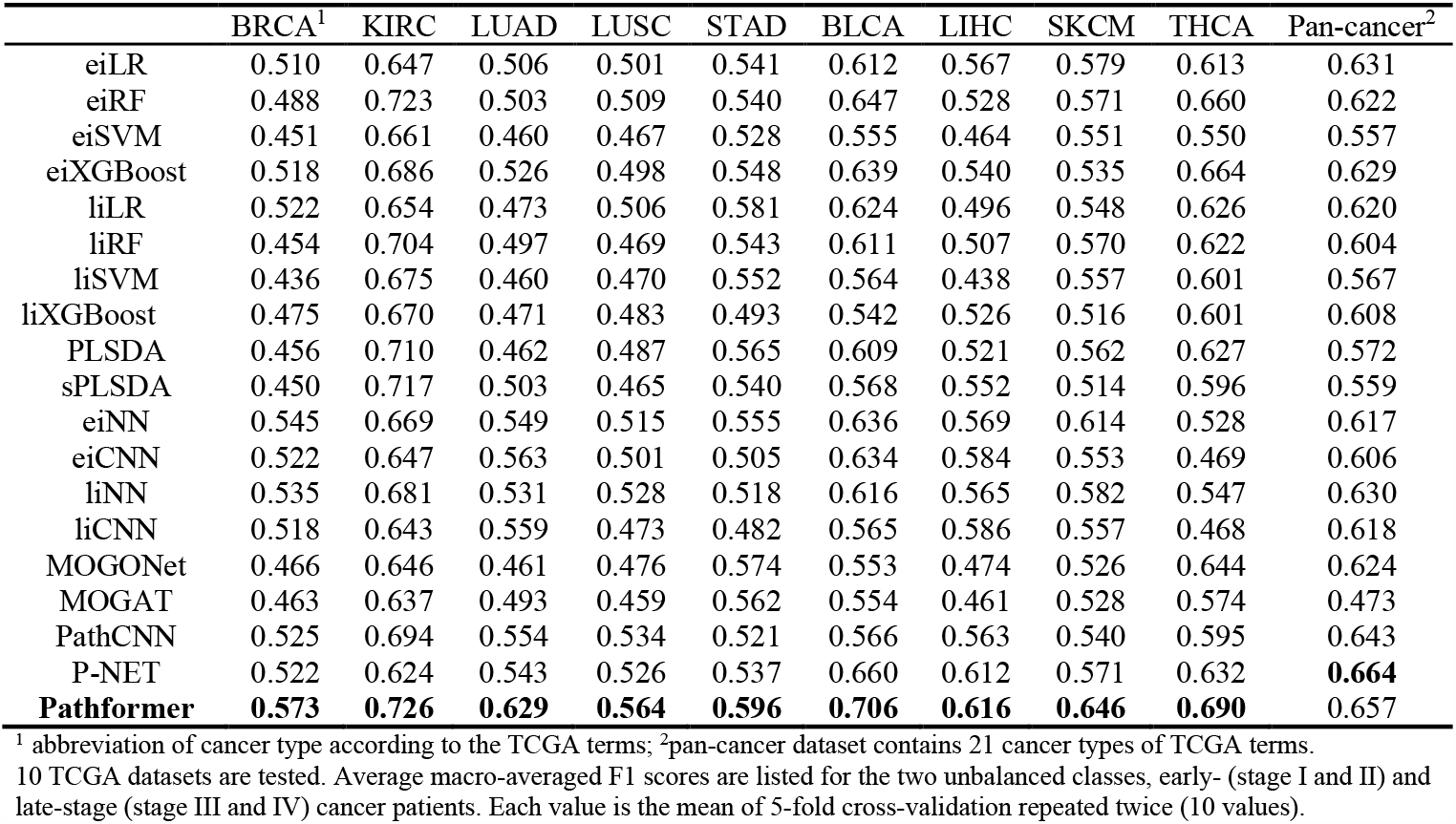
Performance comparison among multiple multi-omics data integration methods for cancer staging.

|  | BRCA<sup>1</sup> | KIRC | LUAD | LUSC | STAD | BLCA | LIHC | SKCM | THCA | Pan-cancer<sup>2</sup> |
| --- | --- | --- | --- | --- | --- | --- | --- | --- | --- | --- |
| eiLR | 0.510 | 0.647 | 0.506 | 0.501 | 0.541 | 0.612 | 0.567 | 0.579 | 0.613 | 0.631 |
| eiRF | 0.488 | 0.723 | 0.503 | 0.509 | 0.540 | 0.647 | 0.528 | 0.571 | 0.660 | 0.622 |
| eiSVM | 0.451 | 0.661 | 0.460 | 0.467 | 0.528 | 0.555 | 0.464 | 0.551 | 0.550 | 0.557 |
| eiXGBoost | 0.518 | 0.686 | 0.526 | 0.498 | 0.548 | 0.639 | 0.540 | 0.535 | 0.664 | 0.629 |
| liLR | 0.522 | 0.654 | 0.473 | 0.506 | 0.581 | 0.624 | 0.496 | 0.548 | 0.626 | 0.620 |
| liRF | 0.454 | 0.704 | 0.497 | 0.469 | 0.543 | 0.611 | 0.507 | 0.570 | 0.622 | 0.604 |
| liSVM | 0.436 | 0.675 | 0.460 | 0.470 | 0.552 | 0.564 | 0.438 | 0.557 | 0.601 | 0.567 |
| liXGBoost | 0.475 | 0.670 | 0.471 | 0.483 | 0.493 | 0.542 | 0.526 | 0.516 | 0.601 | 0.608 |
| PLSDA | 0.456 | 0.710 | 0.462 | 0.487 | 0.565 | 0.609 | 0.521 | 0.562 | 0.627 | 0.572 |
| sPLSDA | 0.450 | 0.717 | 0.503 | 0.465 | 0.540 | 0.568 | 0.552 | 0.514 | 0.596 | 0.559 |
| eiNN | 0.545 | 0.669 | 0.549 | 0.515 | 0.555 | 0.636 | 0.569 | 0.614 | 0.528 | 0.617 |
| eiCNN | 0.522 | 0.647 | 0.563 | 0.501 | 0.505 | 0.634 | 0.584 | 0.553 | 0.469 | 0.606 |
| liNN | 0.535 | 0.681 | 0.531 | 0.528 | 0.518 | 0.616 | 0.565 | 0.582 | 0.547 | 0.630 |
| liCNN | 0.518 | 0.643 | 0.559 | 0.473 | 0.482 | 0.565 | 0.586 | 0.557 | 0.468 | 0.618 |
| MOGONet | 0.466 | 0.646 | 0.461 | 0.476 | 0.574 | 0.553 | 0.474 | 0.526 | 0.644 | 0.624 |
| MOGAT | 0.463 | 0.637 | 0.493 | 0.459 | 0.562 | 0.554 | 0.461 | 0.528 | 0.574 | 0.473 |
| PathCNN | 0.525 | 0.694 | 0.554 | 0.534 | 0.521 | 0.566 | 0.563 | 0.540 | 0.595 | 0.643 |
| P-NET | 0.522 | 0.624 | 0.543 | 0.526 | 0.537 | 0.660 | 0.612 | 0.571 | 0.632 | **0.664** |
| **Pathformer** | **0.573** | **0.726** | **0.629** | **0.564** | **0.596** | **0.706** | **0.616** | **0.646** | **0.690** | 0.657 |

<sup>1</sup> abbreviation of cancer type according to the TCGA terms; <sup>2</sup>pan-cancer dataset contains 21 cancer types of TCGA terms.
10 TCGA datasets are tested. Average macro-averaged F1 scores are listed for the two unbalanced classes, early- (stage I and II) and late-stage (stage III and IV) cancer patients. Each value is the mean of 5-fold cross-validation repeated twice (10 values).

In general, Pathformer significantly performed better than the other 18 integration methods in terms of F1score_macro score (**Fig. 2a**) and cross-validation variances (**Fig. 2b**). In cancer low- and high-risk survival classification tasks, comparing to the other eight deep learning methods (Type III), Pathformer’s F1score_marco showed average improvements between 6.3% and 14.6%. When comparing to eiXGBoost, which performed best in the conventiaonl machine learning methods (Types I and II), Pathformer’s F1score_marco showed an average improvement of 8.3%. In early- and late-stage classification tasks, comparing to the deep learning methods (Type III), Pathformer’s F1score_marco showed average improvements between 5.1% and 12%. Compared to eiXGBoost, Pathformer’s F1score_marco showed an average improvement of 6.2%. Moreover, Pathformer also showed good generalization ability in terms of stability (**Fig. 2b**).

The detailed performance comparisons of Pathformer and other integration methods for different cancer types are shown in **Tables 1-2** and **Supplementary Table 3**. In survival classifications, Pathformer achieved the highest F1score_macro and F1score_weighted in all the 10 datasets, and the highest AUC in 7 of 10 datasets. In stage classifications, Pathformer achieved the highest F1score_macro in 9 of 10 datasets, the highest F1score_weighted in 8 of 10 datasets, and the highest AUC in 6 of 10 datasets.

### Ablation analysis of Pathformer

We used ablation analysis to evaluate the contributions of different input modalities and calculation modules in the Pathformer model, based on 9 datasets for cancer survival prediction (**Fig. 3**) and 9 datasets for cancer staging (**Supplementary Fig. 5**). The pan-cancer dataset of the 10 datasets was not used here. First, we evaluated different input modalities, including RNA expression, DNA methylation, DNA CNV, and a combination thereof. By comparing the performances of seven models on cancer survival risk classification, we discovered that the model with all three modalities as input achieved the best performance, followed by RNA expression and DNA CNV combined model and DNA methylation-only model (**Fig. 3a**). Furthermore, we observed that the performances of models with single modality can vary greatly between datasets. For example, DNA methylation-only model performed better than RNA expression-only and DNA CNV-only in the LUSC, LIHC, and LGG datasets, but the opposite performances were observed in the LUAD and BLCA datasets. Ablation analysis of different modalities on cancer stage classification showed similar results (**Supplementary Fig. 5a**). These findings underscore the distinct behaviors of different modalities in different cancer types, highlighting the necessity of multi-modal data integration in various cancer stage and survival risk classification tasks.

**Figure 3.**
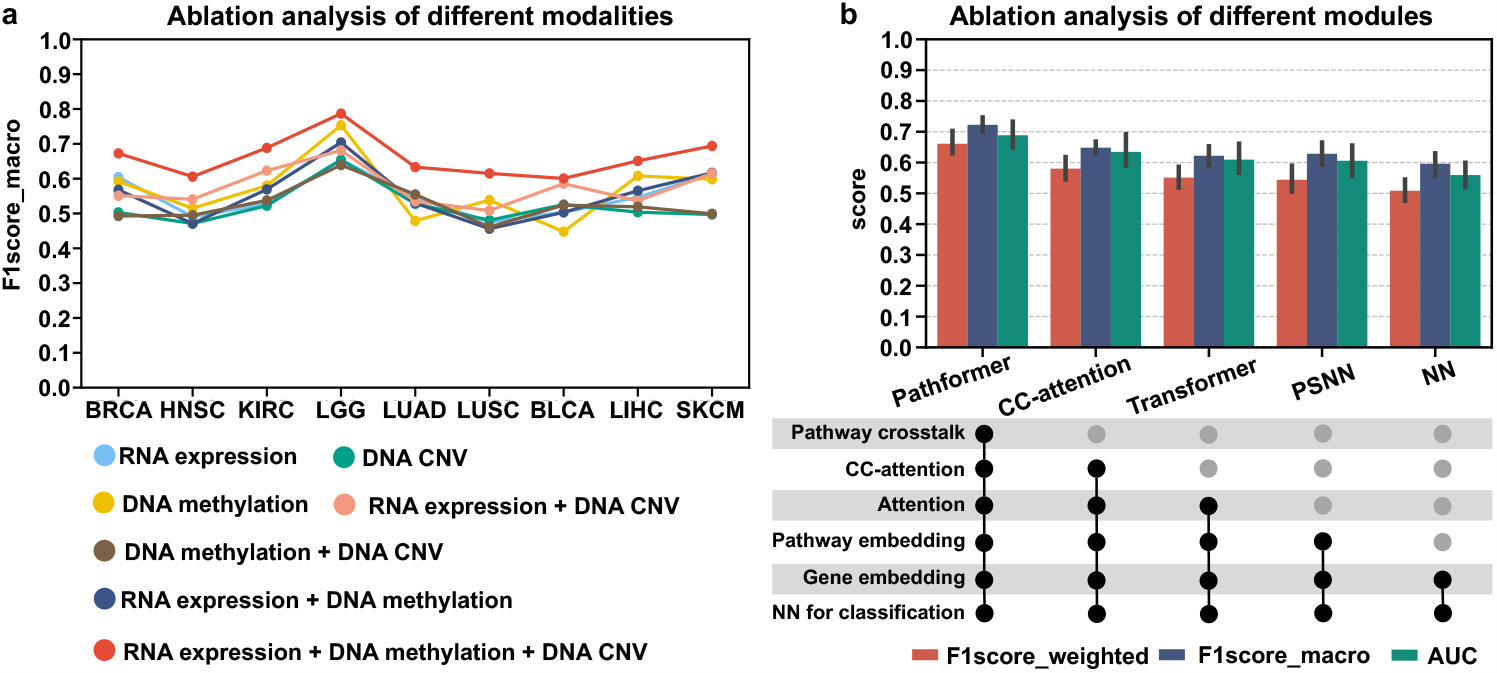
Ablation analysis of Pathformer for different input modalities and different calculation modules. **a**. Different types of input modalities (omics data types) were used as input for TCGA cancer low- and high-risk survival classification. **B**. Ablation analysis of different calculation modules in Pathformer. Error bars are from 2 times 5-fold cross-validation across 9 datasets, representing 95% confidence intervals. CC-attention, Pathformer without pathway crosstalk network bias; Transformer, only based on normal attention and pathway embedding; PSNN, only classification module with pathway embedding; NN, only classification module with gene embedding.

Next, we evaluated the essentialities of different calculation modules in Pathformer. We developed 4 models, namely CC-attention, Transformer, PSNN, and NN, which successively remove one to multiple modules of Pathformer. The “CC-attention” model is Pathformer without pathway crosstalk network bias. The “Transformer” model is Pathformer without pathway crosstalk network bias and row-attention, using only normal attention mechanism and pathway embeddings. The “PSNN” model directly uses classification module with pathway embedding as input. The “NN” model directly uses classification module with gene embedding as input. As shown in **Fig. 3b** and **Supplementary Fig. 5b**, the complete Pathformer achieved the best classification performance, while the performance of CC-Attention, Transformer, PSNN, and NN decreased successively. This indicated that pathway crosstalk network, attention mechanism, and pathway embedding are all integral components of Pathformer. In particular, CC-attention exhibited significantly poorer classification performance compared to Pathformer, providing strong evidence for the necessity of incorporating pathway crosstalk in Pathformer.

### Biological significance revealed by Pathformer for breast cancer prognosis using tissue data

To further understand the decision-making process of Pathformer and validate the reliability of its biological interpretability, we showed a case study on breast cancer survival risk classification. We demonstrated that Pathformer can use attention weights and SHAP values to reveal modalities, pathways, and genes related to breast cancer prognosis, which align with biological knowledges (**Fig. 4**).

**Figure 4.**
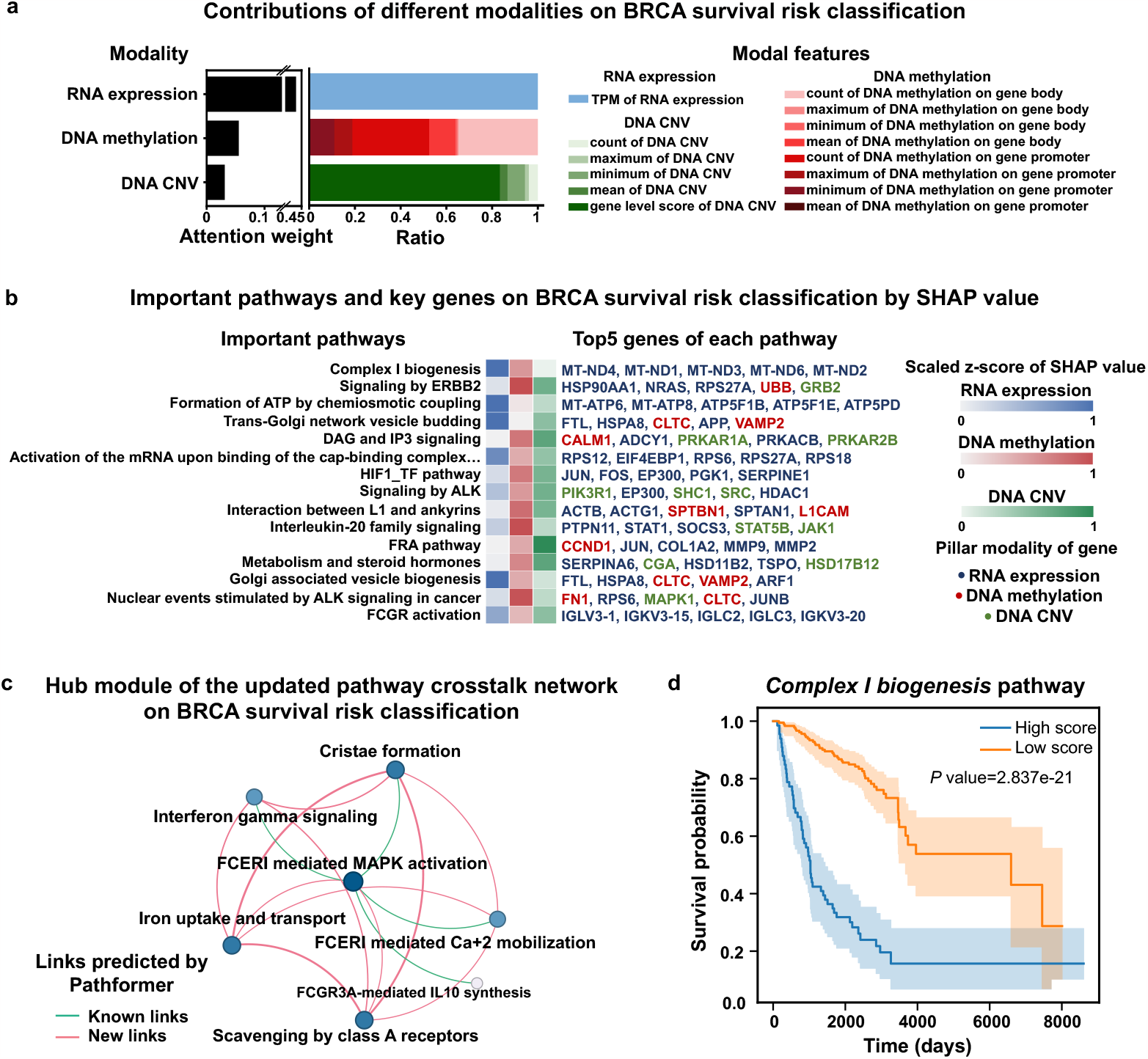
Biological interpretation of the breast cancer survival data using Pathformer. **a**. Contributions of different modalities for breast cancer (BRCA) survival risk classification calculated by attention weights (averaging attention maps of row-attention). **b**. Important pathways and their key genes with top SHapley Additive exPlanations (SHAP) values for BRCA survival risk classification. Among the key genes, different colors represent different pillar modalities of the genes. **c**. A hub module of the updated pathway crosstalk network for BRCA survival risk classification. Color depth and size of node represents the degree of node. Line thickness represents the weight of edge. All links are predicted by Pathformer, where known links are reported by the initial crosstalk network and new links are new predictions. **d**. Keplan-Meier curves of the most active pathway selected identified by Pathformer. P-value calculated through Log-Rank test.

First, at the omics and modality level, we visualized the contributions of different modalities for breast cancer survival risk classification by the attention weights (**Fig. 4a**). The contribution of transcriptomic data was greatest in breast cancer prognostic prediction, which is consistent with the results of ablation analysis (**Fig. 3a**) and findings from other literatures^31,32^. Combining with the results of stage classification using breast cancer tissue multi-modal data (**Supplementary Figs. 6a**), we observed that the contributions of the three modalities in both tasks were similar, but the contributions of different features for DNA methylation in breast cancer prognosis classification were comparable. In other words, the contributions of various features in the same modality were also different for different classification tasks. These findings further validated the necessity of multi-modal integration and biological multi-modal embedding process.

Next, at the pathway and gene levels, we identified key pathways with top 15 SHAP values and key genes with top 5 SHAP values for each pathway in breast cancer survival risk classification (**Fig. 4b**). Then, we presented a hub module of the updated pathway crosstalk network (**Fig. 4c**). Here, *complex I biogenesis* pathway, which participates in cancer cell proliferation and metastasis^33^, was identified as the most critical pathway in the classification and a key node in the hub module of the updated pathway crosstalk network. Five mitochondrial genes (MT-ND4, MT-ND1, MT-ND3, MT-ND6, and MT-ND2) were identified as key genes of the *complex I biogenesis* pathway in breast cancer prognostic classification by Pathformer, which is consistent with the literatures^34^. In addition, compared to stage classification, *Anaplastic Lymphoma Kinase (ALK) signaling* pathway is specific to breast cancer prognosis classification. ALK promotes cell proliferation by inhibiting apoptosis^35^. Overexpression, deletion, and copy number variations of ALK are associated with different breast cancer subtypes^36-38^, especially triple negative breast cancer with poor prognosis^39^. Furthermore, through the hub module of the updated pathway crosstalk network (**Fig. 4c**), we discovered a series of regulatory relationships related to immune response, centered around *FCERI-mediated MAPK activation* pathway^38^, which are crucial to prognosis classification. These pathways impact the tumor immune response by influencing the energy supply^40^, activation^41^, and signal transduction^42^ of immune cells.

Subsequently, to facilitate a more intuitive understanding of the impact of active pathways identified by Pathformer on breast cancer survival risk classification, we depicted survival curves comparing patients with high and low scores in active pathways (**Fig. 4d** and **Supplementary Fig. 7**). The pathway score for each sample was obtained by averaging across different dimensions of pathway embedding. Log-rank tests indicated that most active pathways identified by Pathformer, like *complex I biogenesis* pathway, significantly influenced patient survival.

### Performance of Pathformer for the noninvasive diagnosis of cancer using liquid biopsy data

In clinical practice, cancer diagnosis involves not only using tissue data for cancer staging but also using liquid biopsy data (i.e., plasma) for noninvasive early detection and screen. The later has even greater clinical significance because early detection substantially increases five-year survival rate of cancer patients. For instance, colon cancer’s 5-year survivals were reported as 93.2% for stage I, and only 8.1% for stage IV^43^. Therefore, we applied Pathformer to liquid biopsy data, aiming to classify cancer patients from healthy controls. We curated two types of cell-free RNA sequencing (cfRNA-seq) data derived from plasma and platelet, respectively. We then calculated seven RNA-level modalities as Pathformer’s multi-modal input, including RNA expression, RNA splicing, RNA editing, RNA alternative promoter (RNA alt. promoter), RNA allele-specific expression (RNA ASE), RNA single nucleotide variations (RNA SNV), and chimeric RNA (**Supplementary Note 5**). Because these seven modalities of RNA may have information redundancy, we selected the best modality combination based on 2 times 5-fold cross validations (**Supplementary Note 8**). The results showed that the plasma data with 7 modalities and the platelet data with 3 modalities obtain the best performances (AUCs > 0.9) (**Table 3** and **Supplementary Fig. 9**). Because cancer screen usually requires high specificity, we particularly report sensitivities on 99% specificity in **Table 3**, all of which are above 45%. It is worth noting that the sensitivity is still above 45% on 99% specificity in the plasma data even for the early-stage cancer patients, showing its potential for early cancer diagnosis. In addition, we showed that Pathformer’s performance was superior to the other representative integration methods using the liquid biopsy data (**Supplementary Table 4**).

**Table 3.** Cancer detection performance of Pathformer based on the cell-free RNA liquid biopsy data.

| Dataset | macro-averaged<br>F1 score | weighted-averaged<br>F1 score | AUC | Sensitivity<br>(99% specificity) |
| --- | --- | --- | --- | --- |
| Plasma<sup>1</sup> | 0.843 | 0.877 | 0.914 | 0.488 |
| Plasma (early stage<sup>2</sup>) | 0.853 | 0.869 | 0.916 | 0.479 |
| Platelet<sup>3</sup> | 0.889 | 0.903 | 0.938 | 0.481 |

<sup>1</sup>all cancer stages from I to IV; <sup>2</sup>cancer stage I and stage II; <sup>3</sup>Stage information not available.
All types of cancer patients are used as positives; the healthy controls are used as negatives. Each value is the mean of 5-fold cross-validation repeated twice (10 values).

### Biological significance revealed by Pathformer in cancer patient’s blood

Based on the above analysis, we also gained new insight into the deregulated alterations in plasma through Pathformer’s biological interpretability module. We revealed three interesting pathways deregulated in cancer patients’ blood. The first one is *binding and uptake of ligands* (e.g., oxidized low-density lipoprotein, oxLDL) *by scavenger receptors* pathway, which was identified as the most active pathway in the plasma data (**Fig. 5a**). It is well established that scavenger receptors play a crucial role in cancer prognosis and carcinogenesis by promoting the degradation of harmful substances and accelerating the immune response^44^. Scavenger receptors are also closely related to the transport process of vesicles, such as stabilin-1^45^. Meanwhile, HBB, HBA1, HBA2, FTH1, HSP90AA1 were identified as key genes in this pathway. HBB, as a known biomarker, has been demonstrated to be downregulated in gastric cancer blood transcriptomics^46^. HSP90AA1 is also a potential biomarker for various cancers^47^, especially in the blood^48^.

**Figure 5.**
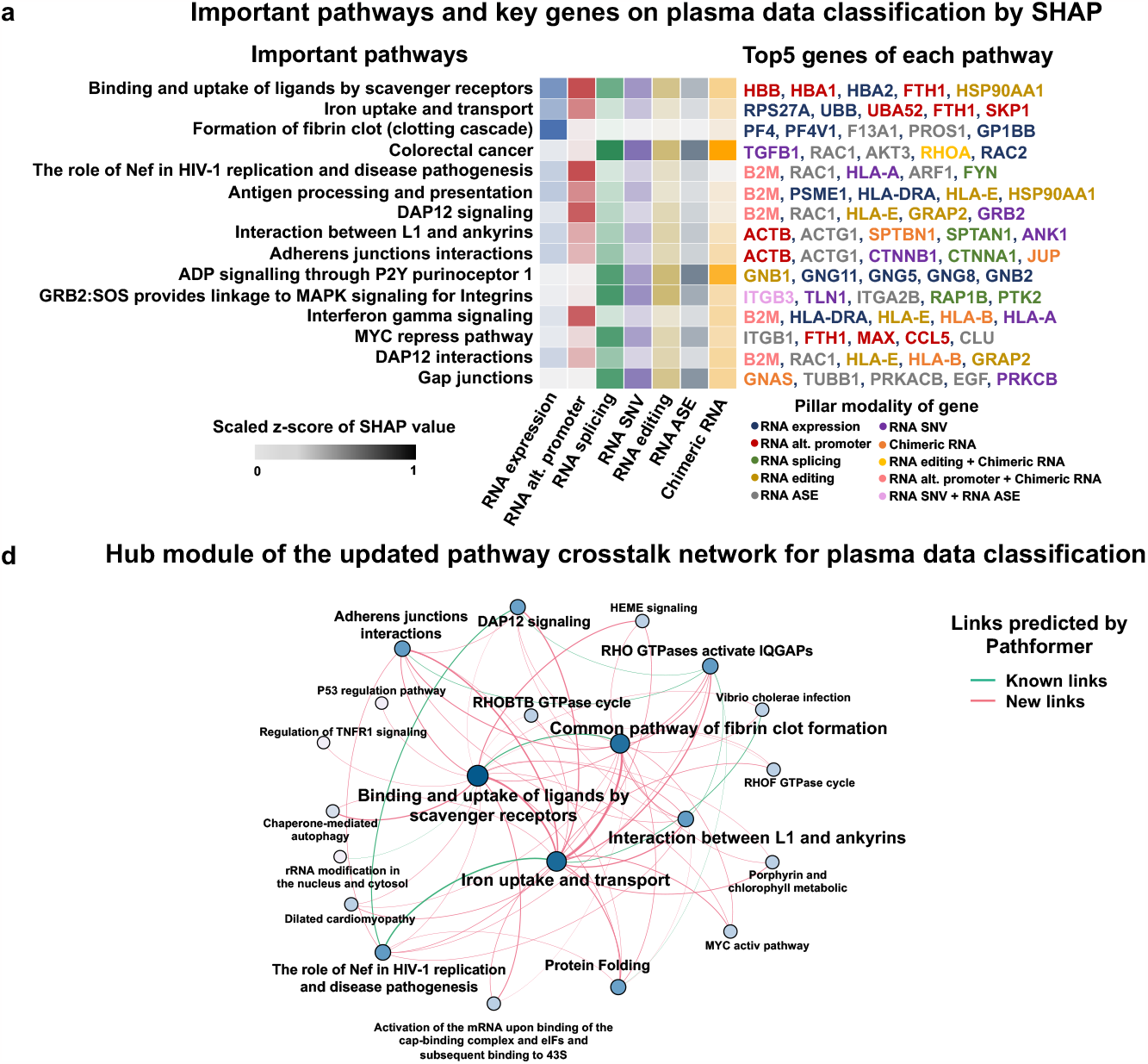
Biological interpretation of the cancer patients’ plasma data using Pathformer. **a**. Important pathways and their key genes revealed by Pathformer in the plasma cell free RNA-seq data when classifying cancer patients from healthy controls. The pathways and their key genes were selected with top SHAP values. Among the key genes, different colors represent different pillar modalities (e.g., RNA expression, RNA editing, etc) of the genes. **b**. Hub modules of pathway crosstalk network are shown for plasma cell free RNA-seq data. Color depth and size of node represent the degree of node. Line thickness represents the weight of edge. All links are predicted by Pathformer, where known links are reported by the initial crosstalk network and new links are new predictions.

The other two interesting pathways, *DAP12 signaling* pathway and *DAP12 interactions* pathway, were revealed in both plasma and platelet data (**Fig. 5a and Supplementary Fig. 10a**). DAP12 triggers natural killer cell immune responses against certain tumor cells^49^, which is regulated by platelet^50^. Among the top 5 key genes of DAP12 related pathway, B2M has been reported as a serum protein encoding gene and a widely recognized tumor biomarker^51^; HLA-E and HLA-B were reported as cancer biomarkers in tissue and plasma^52,53^.

Furthermore, Pathformer provides insight into the interplay between various biological processes and their impact on cancer progression by updating pathway crosstalk network (**Fig. 5b**). In the plasma data, the link between *binding and uptake of ligands by scavenger receptors* pathway and *iron uptake and transport* pathway was a novel addition to the updated network. These two pathways lack the crosstalk relationship in the original network before training, but there are literature reports that SCARA5 (member of scavenger receptor class A) can function as a ferritin receptor^54^. The crosstalk between two pathways was amplified by Pathformer in plasma dataset, probably because they were important for classification and shared the same key gene, FTH1 (gene encoding the heavy subunit of ferritin), one of two intersecting genes between the two pathways. In summary, Pathformer’s updated pathway crosstalk network visualizes the information flow between pathways related to cancer classification task in the liquid biopsy data, providing novel insight into the crosstalk of biological pathways in cancer patients’ plasma.

## Conclusion and Discussion

Pathformer successfully applied a Transformer model to the integrate multi-modal data for cancer diagnosis and prognosis. Particularly, it introduced a novel biological embedding method based on the compacted multi-modal vectors (**Fig. 1b**). Moreover, it utilized the criss-cross attention mechanism of Transformer to capture crosstalk between biological pathways and regulation between modalities (i.e., different omics).

### Clinical Applications of Pathformer

Pathformer can be applied to various classification tasks in disease diagnosis and prognosis, such as early detection, cancer staging, and survival prediction. Its prediction performance and biological interpretability were demonstrated through substantial benchmark and case studies focusing on cancer prognosis and noninvasive diagnosis. Moreover, this framework is adaptable for the diagnosis and prognosis of other complex diseases, like autoimmune disease, neurodegenerative diseases, etc.

### Potential targets revealed in cancer patients’ blood

Particularly, we revealed the scavenger receptor related pathway and DAP12 related pathway in cancer patients’ blood, which are associated with extracellular vesicle transport^45^ and immune response^49^, respectively. We even found a new cancer-related pathway crosstalk in blood, which is between *binding and uptake of ligands by scavenger receptors* pathway and *iron uptake and transport* pathway. These results provide candidate targets for the mechanism study of cancer microenvironment and immune system, and even new targets for cancer treatment.

### Limitations of Pathformer and future directions

Pathformer used genes involved in pathways from four public databases, all of which consist of protein-coding genes. However, a substantial body of literature has reported that noncoding RNAs are also crucial in cancer prognosis and diagnosis^55^. Therefore, incorporating noncoding RNAs and their related functional pathways into Pathformer would be promising for future work. In addition, we used the multi-modal features derived from cfRNA-seq only in the application of liquid biopsy, because the published cell-free multi-omics datasets^28^ are usually too small to be train-and-tested. Another flaw of Pathformer is the computing memory issue. Pathway embedding of Pathformer has prevented memory overflow of Transformer module caused by long inputs. However, when adding more pathways or gene sets (e.g., transcription factors), Pathformer still faces the issue of memory overflow. In the future work, we may introduce linear attention to further improve computational speed.

## Supporting information

Supplementary Table 1.Detail of gene embedding method for different datasets

Supplementary Table 2.Results of optimal hyperparameter combinations of TCGA datasets

Supplementary Table 3.Performances of multiple methods on TCGA datasets

Supplementary Table 4.Comparison results of Pathformer and other representative methods on liquid biopsy datasets

## Declarations

### Data availability

All datasets used in this study are publicly available for academic research usages. The details of usage are also fully illustrated in Methods and Supplementary Notes.

### Code availability

Source code for data preprocessing and model training is freely available at Github (https://github.com/lulab/Pathformer) with detailed instructions. Source code for comparing the other methods is also included.

### Consent for publication

All authors have approved the manuscript and agree with the publication.

### Funding and Acknowledgements

This work is supported by National Natural Science Foundation of China (32170671, 8237061277), Tsinghua University Guoqiang Institute Grant (2021GQG1020), Tsinghua University Initiative Scientific Research Program of Precision Medicine (2022ZLA003), Bioinformatics Platform of National Center for Protein Sciences (Beijing) (2021-NCPSB-005). This study was also supported by Bayer Micro-funding, Bio-Computing Platform of Tsinghua University Branch of China National Center for Protein Sciences.

