## Supplementary information and Notes for "Pathformer: a biological pathway informed Transformer integrating multi-omics data for disease diagnosis and prognosis"

### 39 Supplementary Figures

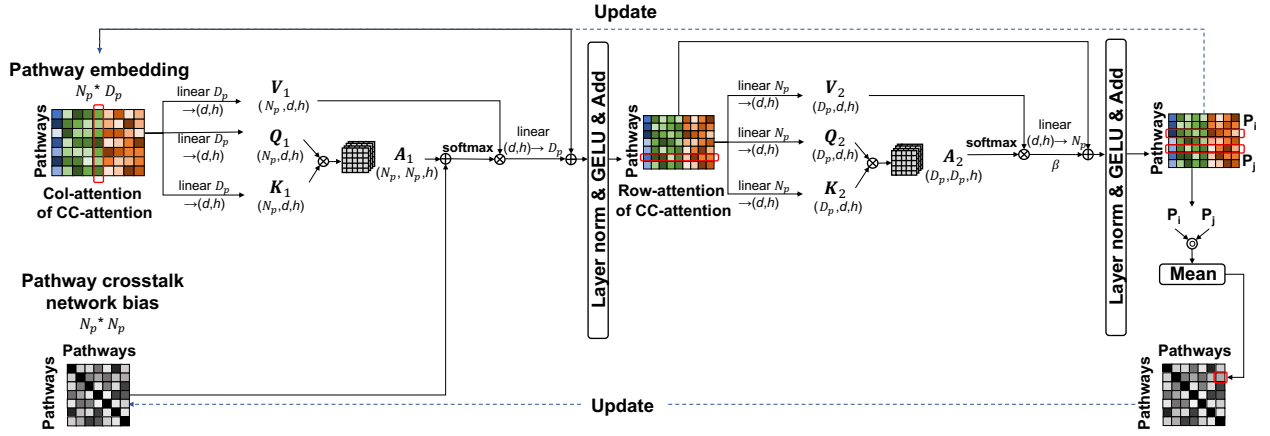

41 **Supplementary figure 1: A block of Transformer module with pathway crosstalk network bias.**

42 The pathway embedding matrix is used as input and the pathway crosstalk network matrix is used as bias. $N_p$,
43 number of pathways; $D_p$, dimensionality of pathway embedding; $h$, number of attention heads; $d$, attention
44 dimension; $V_1, K_1, Q_1, A_1$: value, key, query and attention map of col-attention; $V_2, K_2, Q_2, A_2$: value, key, query
45 and attention map of row-attention; $+$, element-wise addition; ×, matrix multiplication; $\odot$, matrix dot product; $\beta$,
46 constant coefficient for row-attention.

### Benchmark Datasets

#### TCGA multi-omics data

##### Early- and late-stage classification

**BRCA** (544 early-stage, 208 late-stage),
**KIRC** (184 early-stage, 130 late-stage),
**LUAD** (354 early-stage, 92 late-stage),
**LUSC** (301 early-stage, 60 late-stage),
**STAD** (152 early-stage, 174 late-stage),
**BLCA** (131 early-stage, 268 late-stage),
**LIHC** (256 early-stage, 84 late-stage),
**SKCM** (152 early-stage, 193 late-stage),
**THCA** (332 early-stage, 113 late-stage),
**Pan-cancer** (3440 early-stage, 2170 late-stage)

##### Low- and high-risk survival classification

**BRCA** (178 low-risk, 69 high-risk),
**HNSC** (50 low-risk, 190 high-risk),
**KIRC** (93 low-risk, 89 high-risk),
**LGG** (64 low-risk, 98 high-risk),
**LUAD** (48 low-risk, 146 high-risk),
**LUSC** (61 low-risk, 128 high-risk),
**BLCA** (47 low-risk, 169 high-risk),
**LIHC** (40 low-risk, 119 high-risk),
**SKCM** (151 low-risk, 143 high-risk),
**Pan-cancer** (1388 low-risk, 2059 high-risk)

#### Supplementary figure 2: Benchmark datasets.

We collected cancer datasets from TCGA covering two types of tasks: cancer early- and late-stage classification and cancer high- and low-risk survival classification. For cancer early- and late- stage classification, we defined stage I and stage II as the early-stage and stage III as the late-stage. For cancer low- and high- survival risk classification, we defined samples with survival time greater than 1825 days as low-risk and those less than 1825 days as high-risk. Each dataset name is an abbreviation of cancer type according to the TCGA terms. The pan-cancer dataset contains 33 cancer types of TCGA terms. More details are in Supplementary Note 5.

**a BRCA early- and late- stage classification**

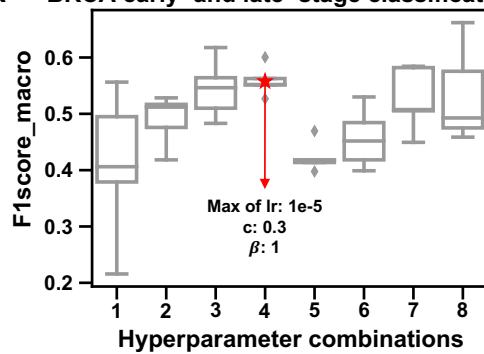

**b BRCA high- and low- risk classification**

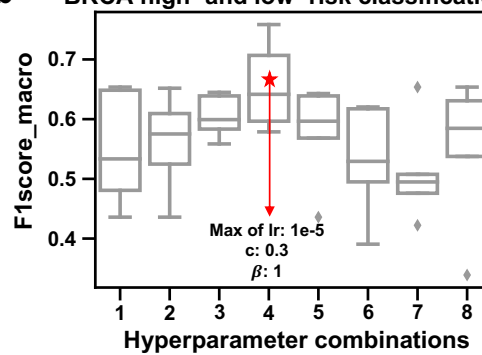

**Supplementary figure 3: Optimal combination of hyperparameters.**

Optimal combination of hyperparameters in one experiment for (a) BRCA early- and late- stage classification and (b) BRCA high- and low- risk survival classification, demonstrating how to optimize parameters. First, we gave some hyperparameters of Pathformer based on prior experience, such as the number of blocks and the number of multi-head self-attention. Subsequently, we aimed to determine the optimal value of 8 combinations consisting of maximum of learning rate, dropout probability of classification ($c$), and constant coefficient for row-attention ($\beta$). Then, we performed 5-fold cross-validation on the training set for hyperparameter optimization. We used the F1score\_macro as the selection criterion and used grid search to find the optimal combination of hyperparameters.

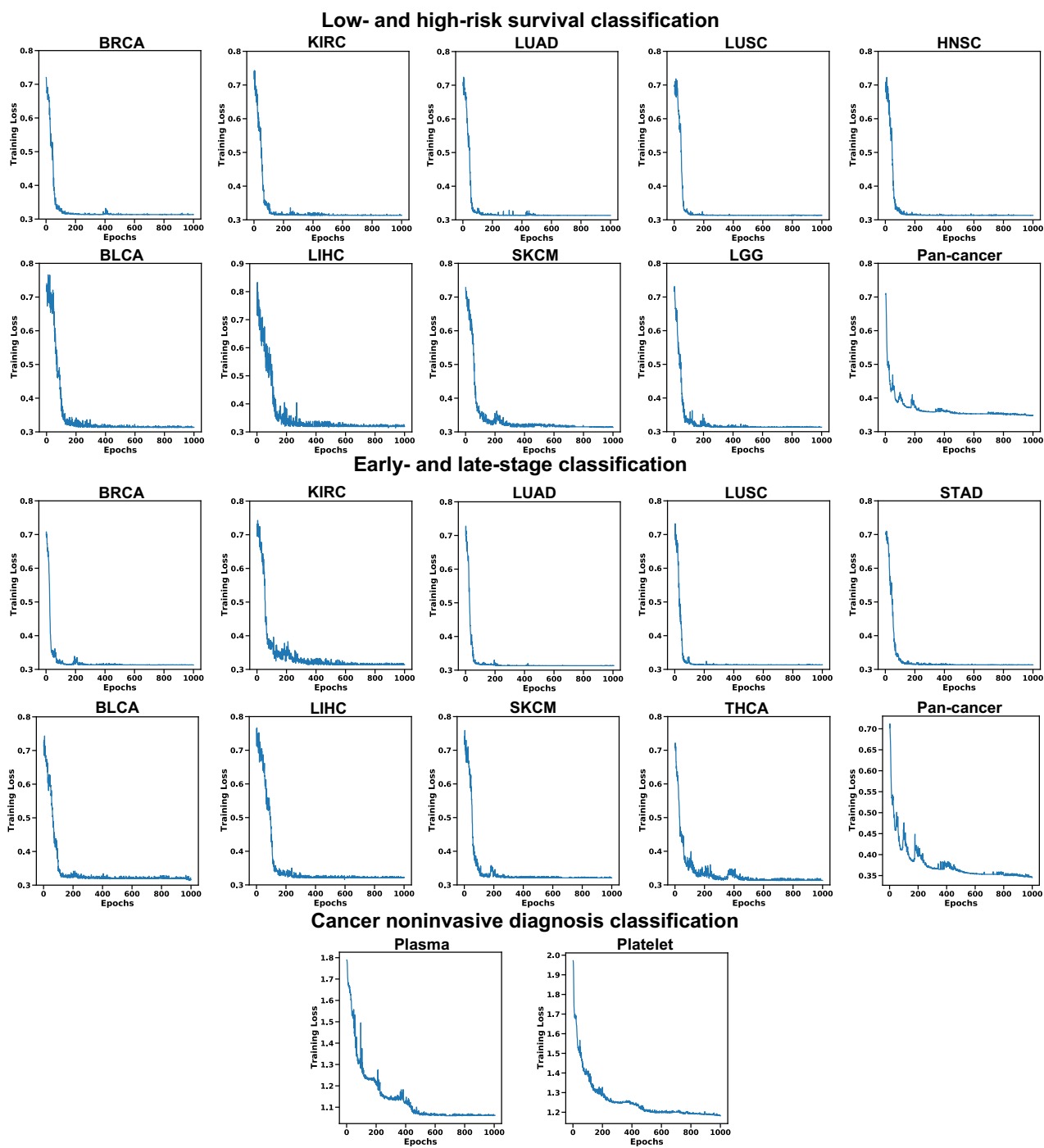

### **Supplementary figure 4: Convergence analysis of Pathformer.**

We depicted the training loss in relation to the number of epochs on multiple TCGA datasets and the liquid biopsy datasets, verifying the convergence of Pathformer. The total loss for each dataset decreases rapidly as the iterations progress, ultimately converging to a stable state. Furthermore, the specific number of training epochs for Pathformer on each dataset is determined by the stopping criteria of the early stopping strategy (for more details, refer to Supplementary Note 6).

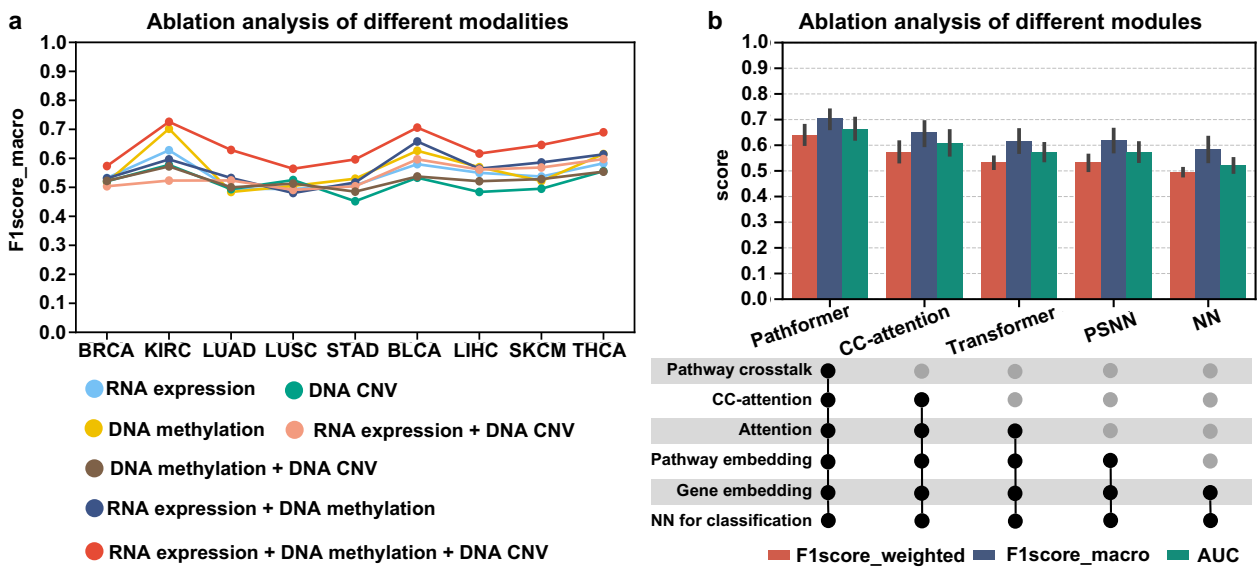

**Supplementary figure 5: Ablation analysis of Pathformer for the classification of early- and** **late-stage cancer patients.**

a. Different types of input modalities (omics data types) were used as input for TCGA cancer early- and late-stage classification. B. Ablation analysis of different calculation modules in Pathformer. Error bars are from 2 times 5-fold cross-validation across 9 datasets, representing 95% confidence intervals. CC-attention, Pathformer without pathway crosstalk network bias; Transformer, only based on normal attention and pathway embedding; PSNN, only classification module with pathway embedding; NN, only classification module with gene embedding.

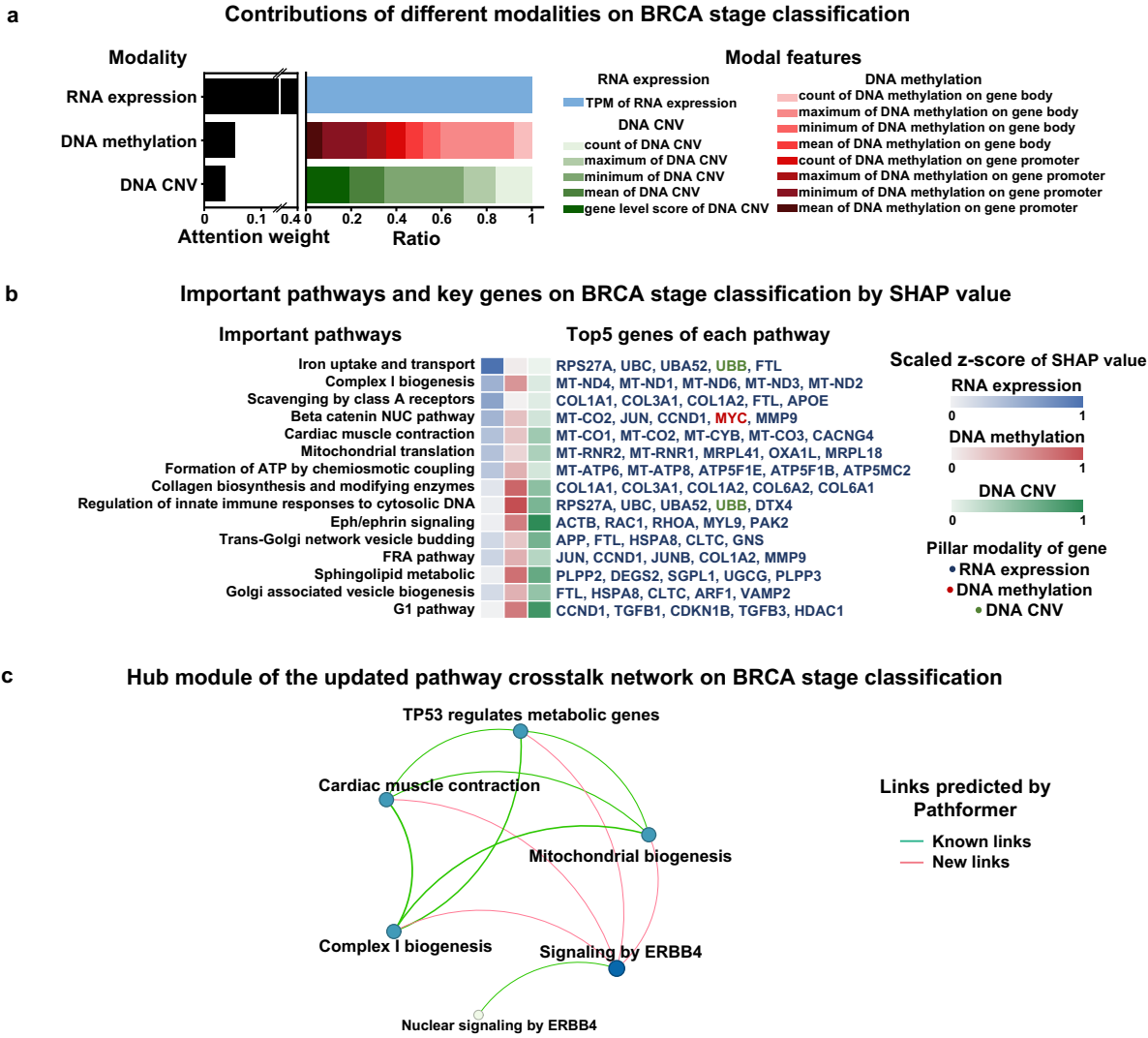

**Supplementary figure 6: BRCA early- and late- stage classification related modalities,** **pathways and genes revealed by Pathformer.**

**a.** Contributions of different modalities for BRCA early- and late- stage classification calculated by attention weights. **b.** Important pathways and their key genes with top SHapley Additive exPlanations (SHAP) values. Among the key genes, different colors represent different pillar modalities of the genes. **c.** A hub module of the updated pathway crosstalk network for BRCA early- and late- stage classification. Color depth and size of node represents the degree of node. Line thickness represents the weight of edge. All links are predicted by Pathformer, where known links are reported by the initial crosstalk network and new links are new predictions. In breast cancer early- and late-stage classification, *iron uptake and transport* pathway had the greatest impact. Supportively, the transport and storage of iron in cells are known to play a key role in carcinogenesis, cell proliferation, and the development of breast cancer<sup>1</sup>.

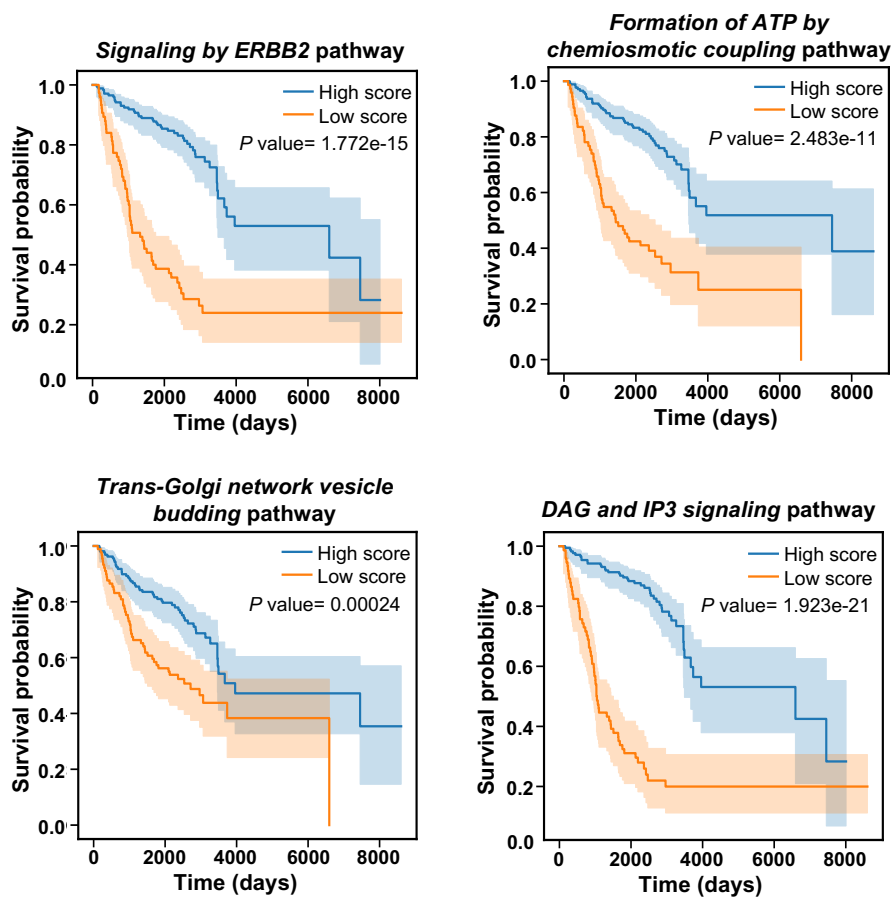

**Supplementary figure 7: Kaplan-Meier curves of active pathway selected identified by** **Pathformer on BRCA survival risk classification.**

Kaplan-Meier curves is depicted the hierarchical relationship between patients with high and low scores in active pathways. The pathway score for each sample was obtained by averaging across different dimensions of pathway embedding. P-value calculated through Log-Rank test.

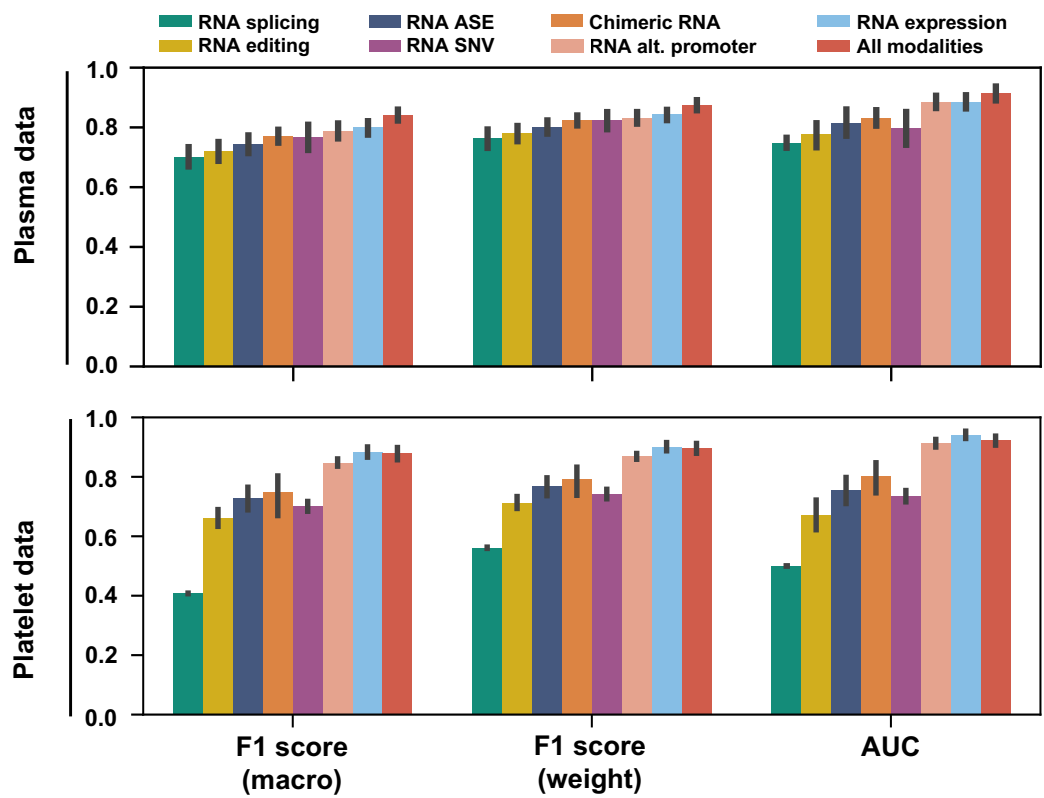

**Supplementary figure 8: Performances of Pathformer with different modalities on liquid biopsy** **datasets.**

Error bars obtained from 2 times 5-fold cross-validation and represent 95% confidence intervals of various evaluation scores.

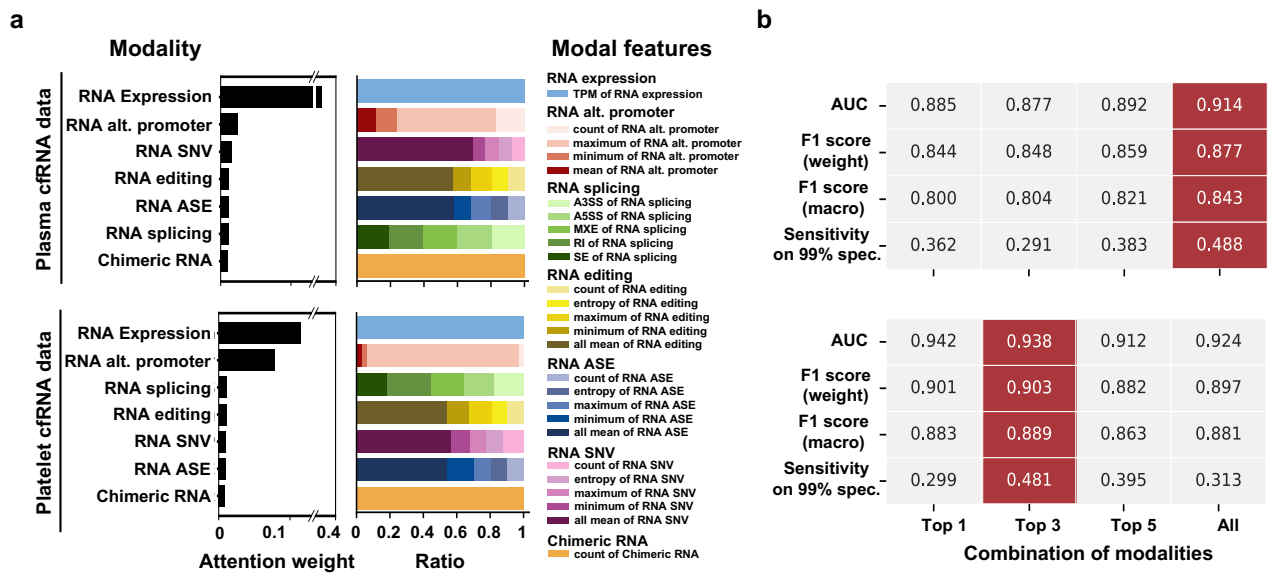

**Supplementary figure 9: Pathformer integrates multi-modal liquid biopsy data for noninvasive cancer diagnosis.**

**a.** Contributions of different input features and their statistical indicators when classifying cancer patients from healthy controls by seven RNA-level modalities on two liquid biopsy datasets (cell free RNA-seq). All mean represents the sum of mean, weighted mean and window weighted mean. Each type of RNA splicing is the sum of all statistical indicators in this type. **b.** Classification performance of different input combinations on two liquid biopsy datasets. Each value is the mean of 2 times 5-fold cross-validation. Top 1, Top 3, and Top 5 respectively represent the top-contributing modality combinations, the top three-contributing modality combinations, and the top five-contributing modality combinations, evaluated by Pathformer on each dataset. For example, in plasma dataset, the top three-contributing modality combinations are RNA expression, RNA alternative promoter, and RNA single nucleotide variations. In platelet dataset, the top three-contributing modality combinations are RNA expression, RNA alternative promoter, and RNA splicing.

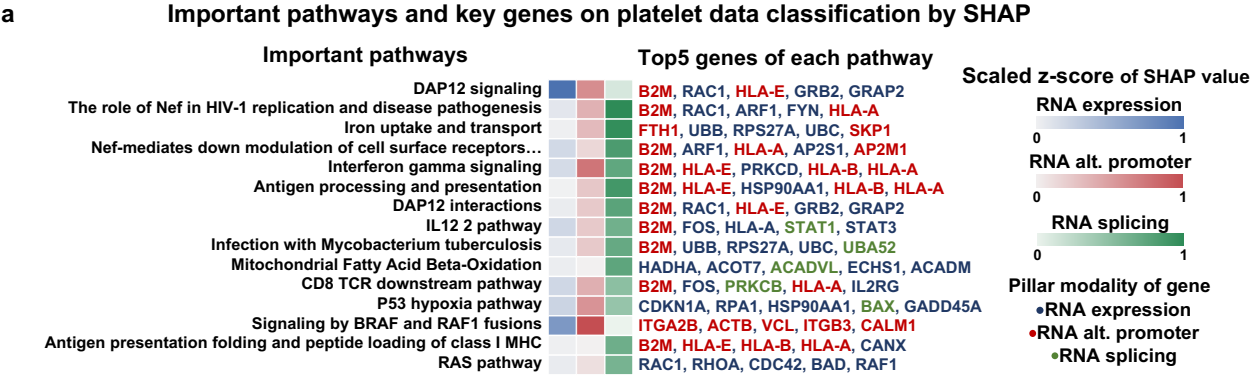

**b** Hub module of the updated pathway crosstalk network for platelet data classification

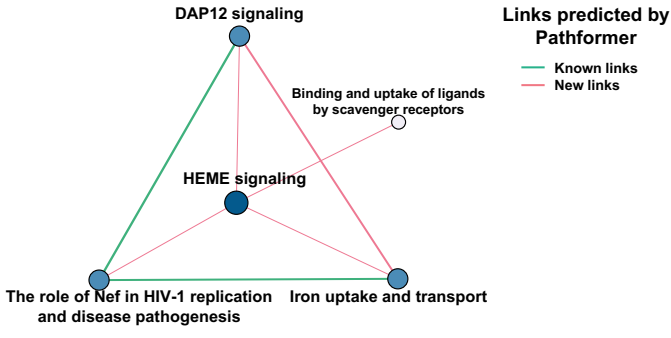

**Supplementary figure 10: Interpretation of cancer patients' platelet data using Pathformer.**

**a.** Important pathways and their key genes revealed by Pathformer in the platelet cell free RNA-seq data when classifying cancer patients from healthy controls. The pathways and their key genes were selected with top SHAP values. Among the key genes, different colors represent different pillar modalities of the genes. **b.** Hub modules of pathway crosstalk network are shown for platelet cell free RNA-seq data. Color depth and size of node represent the degree of node. Line thickness represents the weight of edge. All links are predicted by Pathformer, where known links are reported by the initial crosstalk network and new links are new predictions.

### 134 **Supplementary Notes**

#### 135 **Supplementary Note 1: Pathway crosstalk network calculation**

We used *BinoX*, a classic tool for crosstalk analysis, to calculate the crosstalk relationship of 1,497 pathways and form a pathway crosstalk network. *BinoX* uses the relationship between genes of two pathways in genome-wide functional association networks to calculate the degree of association between two pathways, that is, the crosstalk between two pathways. We used FunCoup v3.0 database<sup>2</sup> to obtain genome-wide functional association networks. Then, we set the cut off for the link weight to 0.75, the number of iterations for the sampling method to 100, and the minimum number of nodes per group to 15 on *BinoX* software.

#### **Supplementary Note 2: Conversion function in gene embedding**

In gene embedding, we use the conversion function $\mathbf{F}_E$, a series of statistical indicator functions, to uniformly convert different modalities into gene level modal features. These statistical indicator functions include gene level score ($f_1$), count ($f_2$), minimum ($f_3$), maximum ($f_4$), mean ($f_5$), entropy ($f_6$), weighted mean in whole gene ($f_7$) and weighted mean in window ($f_8$). The formulas are as follows:

$$147 \quad f_1(\mathbf{M}_i) = [X_{g_1}^{(i)}, \dots, X_{g_{N_g}}^{(i)}] \in \mathbb{R}^{N_g}$$

$$148 \quad f_2(\mathbf{M}_i) = [\text{count}(\mathbf{X}_{ge_1}^{(i)}), \dots, \text{count}(\mathbf{X}_{ge_{N_g}}^{(i)})] \in \mathbb{R}^{N_g}$$

$$149 \quad f_3(\mathbf{M}_i) = [\min(\mathbf{X}_{ge_1}^{(i)}), \dots, \min(\mathbf{X}_{ge_{N_g}}^{(i)})] \in \mathbb{R}^{N_g}$$

$$150 \quad f_4(\mathbf{M}_i) = [\max(\mathbf{X}_{ge_1}^{(i)}), \dots, \max(\mathbf{X}_{ge_{N_g}}^{(i)})] \in \mathbb{R}^{N_g}$$

$$151 \quad f_5(\mathbf{M}_i) = [\text{mean}(\mathbf{X}_{ge_1}^{(i)}), \dots, \text{mean}(\mathbf{X}_{ge_{N_g}}^{(i)})] \in \mathbb{R}^{N_g}$$

$$152 \quad f_6(\mathbf{M}_i) = [\text{entropy}(\mathbf{X}_{ge_1}^{(i)}), \dots, \text{entropy}(\mathbf{X}_{ge_{N_g}}^{(i)})] \in \mathbb{R}^{N_g}$$

$$153 \quad f_7(\mathbf{M}_i) = [\text{weighted\_mean}(\mathbf{X}_{ge_1}^{(i)}), \dots, \text{weighted\_mean}(\mathbf{X}_{ge_{N_g}}^{(i)})] \in \mathbb{R}^{N_g}$$

$$154 \quad \text{weighted\_mean}(\mathbf{X}_{ge_l}^{(i)}) = (mc_1^{(i)(l)} + \dots + mc_{t_l}^{(i)(l)}) / (tc_1^{(i)(l)} + \dots + tc_{t_l}^{(i)(l)}), x_t^{(i)(l)} = mc_t^{(i)(l)} / tc_t^{(i)(l)}$$

$$155 \quad f_8(\mathbf{M}_i) = [[f_7(\mathbf{M}_i^{\text{window1}}), f_7(\mathbf{M}_i^{\text{window2}}), f_7(\mathbf{M}_i^{\text{window3}})]] \in \mathbb{R}^{N_g \times 3}$$

156 , where $X_{g_l}^{(i)}$ is the score of the $i$th modality corresponding to the $l$th gene, $\mathbf{X}_{ge_l}^{(i)} = [x_1^{(i)(l)}, x_2^{(i)(l)}, \dots, x_{t_l}^{(i)(l)}]$ is the
157 event vector mapped to the $l$th gene in the $i$th modality; $t_l$ is the number of events mapped to the $l$th gene; $x_t^{(i)(l)}$ is
158 the score of the $t$th event of the $l$th gene in the $i$th modality; $mc_t^{(i)(l)}$, $tc_t^{(i)(l)}$ are the number of mutated reads and total reads of the $t$th event of the $l$th gene in the $i$th modality; $\mathbf{M}_i^{window1} = [X_{g_1}^{(i)(window1)}, \dots, X_{g_{N_g}}^{(i)(window1)}]$, and $X_{g_l}^{(i)(window1)}$ is 1/3 event vector mapped to the $l$th gene in the $i$th modality. Conversion function of modality $i$ is constructed from distinct statistical indicator functions (more details in **Supplementary Table 1**). When the length of gene embedding $D_g$ is still less than 2 after processing by conversion functions $\mathbf{F}_E$, we use a fully connected neural network layer to transform gene embedding to 32 dimensions.

#### Supplementary Note 3: Classification calculation of Pathformer

Pathformer uses the Transformer module based on criss-cross attention with pathway crosstalk network bias, which has 3 blocks. We used superscripts with parenthesis to represent data at different layers, where $\mathbf{E}_p^{(0)} = \mathbf{E}_p$, $\mathbf{P}^{(0)} = \mathbf{P}$ is the data before entering the first layer. Pathformer is calculated as follows:

$$\begin{aligned} \mathbf{E}_p^{(1)}, \mathbf{P}^{(1)} &= \text{Transformer}(\mathbf{E}_p^{(0)}, \mathbf{P}^{(0)}) \\ \mathbf{E}_p^{(2)}, \mathbf{P}^{(2)} &= \text{Transformer}(\mathbf{E}_p^{(1)}, \mathbf{P}^{(1)}) \\ \mathbf{E}_p^{(3)}, \mathbf{P}^{(3)} &= \text{Transformer}(\mathbf{E}_p^{(2)}, \mathbf{P}^{(2)}) \end{aligned}$$

In order to solve classification tasks, we used a fully connected neural network as the classification module to transform pathway embedding encoded by the Transformer module into the probability for each label. The calculation is as follows:

$$\begin{aligned} \mathbf{L}_1 &= \text{Flatten}(\mathbf{E}_p^{(3)}) \\ \mathbf{Z}_1 &= \text{dropout}_c(\text{RELU}(\mathbf{L}_1 \mathbf{W}_{c1} + \mathbf{B}_{c1})) \\ \mathbf{Z}_2 &= \text{dropout}_c(\text{RELU}(\mathbf{Z}_1 \mathbf{W}_{c2} + \mathbf{B}_{c2})) \\ \mathbf{Z}_3 &= \text{dropout}_c(\text{RELU}(\mathbf{Z}_2 \mathbf{W}_{c3} + \mathbf{B}_{c3})) \\ \hat{\mathbf{Y}} &= \text{softmax}(\mathbf{Z}_3 \mathbf{W}_{out} + \mathbf{B}_{out}) \end{aligned}$$

, where $\mathbf{W}_{c1} \in \mathbb{R}^{(N_p * D_p) \times d_{c1}}$, $\mathbf{W}_{c2} \in \mathbb{R}^{d_{c1} \times d_{c2}}$, $\mathbf{W}_{c3} \in \mathbb{R}^{d_{c2} \times d_{c3}}$, $\mathbf{W}_{out} \in \mathbb{R}^{d_{c3} \times d_{out}}$ are the weight matrices as parameters; $d_{c1} = 300$, $d_{c2} = 200$, $d_{c3} = 100$ are numbers of neurons for three fully connected neural network layers; $d_{out}$ is the number of sample classes in classification tasks; $\mathbf{B}_{c1}$, $\mathbf{B}_{c2}$, $\mathbf{B}_{c3}$ are bias terms; Flatten function is a flatten layer that collapses the input; RELU is an activation function; softmax is a normalized exponential function; $\text{dropout}_c$ is a dropout neural network layer with a probability of $c$; $\hat{\mathbf{Y}}$ is the probability for each label.

##### 184 **Supplementary Note 4: Biological interpretability module**

To comprehend Pathformer's decision-making process, we used averaging attention maps in row-attention to represent the contributions of different modalities, and SHAP value to decipher the important pathways and their key genes. Finally, the hub module of the updated pathway crosstalk network represents the most critical regulatory mechanism in classification.

###### ***Contribution of each modality***

In Pathformer, row-attention is used to facilitate information interaction between different modalities, that is, row-attention map can represent the importance of each modality. According to the trained model, we obtained row-attention maps of 8 heads in 3 blocks for each sample. For the contribution of each modality, we first integrated all matrices of row-attention maps into one matrix by element-wise average. Then, we averaged this average row-attention matrix along with columns as the attention weights of modalities, i.e., the contribution of modalities. The calculation is as follows:

$$196 \quad \mathbf{A}_{aver} = \frac{1}{N} \sum_{n=1}^N \frac{1}{BL} \sum_{b=1}^{BL} \frac{1}{H} \sum_{h=1}^H softmax([[\mathbf{A}_2^{(h)}]^{(b)}]^{(n)})$$

$$197 \quad attention\ weight_i = \frac{1}{D_p} \sum_{j=1}^{D_p} a_{ij}, a_{ij} \text{ is the } i\text{th row and the } j\text{th columns of } \mathbf{A}_{aver}$$

, where $N$ is the number of samples, $BL$ is the number of blocks, $H$ is the number of heads, softmax is a normalized exponential function, and $attention\ weight_i$ is the attention weight of dimension $i$ of pathway embedding.

###### ***Important pathways and their key genes***

We calculated SHAP values of the gene embedding and the pathway embedding encoded by Transformer module corresponding to each sample and each category, denoted as $\mathbf{S}_{gn}^{(j)} \in \mathbb{R}^{D_p}$ and $\mathbf{S}_{pn}^{(j)} \in \mathbb{R}^{D_p}$ respectively. The SHAP values of genes and pathways are calculated as follows:

$$204 \quad SHAP_g = \sum_{j=1}^{d_{out}} \sum_{e=1}^{D_p} \frac{1}{N} \sum_{n=1}^N |s_{gne}^{(j)}|, s_{gne}^{(j)} \in \mathbf{S}_{gi}^{(j)}$$

$$205 \quad SHAP_p = \sum_{j=1}^{d_{out}} \sum_{e=1}^{D_p} \frac{1}{N} \sum_{n=1}^N |s_{pne}^{(j)}|, s_{pne}^{(j)} \in \mathbf{S}_{pi}^{(j)}$$

, where $g = 1, 2, \dots, N_g$ is the $g$th gene, $p = 1, 2, \dots, N_p$ is the $p$th pathway, $n = 1, 2, \dots, N$ is the $n$th sample, $e =$ $1, 2, \dots, D_p$ is dimension $e$ of pathway embedding, and $j = 1, 2, \dots, d_{out}$ is the $j$th category of sample.

In addition, the z-score of SHAP values of different modalities for each pathway and gene can demonstrate modal complementarity at the gene level and the pathway level, described as follows:

$$\text{SHAP}_{gi} = \sum_{j=1}^{d_{out}} \sum_{e=e_1+\dots+e_{i-1}}^{e_i} \frac{1}{N} \sum_{n=1}^N |s_{gne}^{(j)}|, s_{gie}^{(j)} \in \mathbf{S}_{gi}^{(j)}$$

$$\text{SHAP}_{pi} = \sum_{j=1}^{d_{out}} \sum_{e=e_1+\dots+e_{i-1}}^{e_i} \frac{1}{N} \sum_{n=1}^N |s_{pne}^{(j)}|, s_{pie}^{(j)} \in \mathbf{S}_{pi}^{(j)}$$

, where $i = 1, \dots, m$ is the $i$th modality, $e_i$ is the length of gene embedding and pathway embedding for modality $i$.

Finally, pathways with the top 15 SHAP values in the classification task are considered as important pathways.

For each pathway, genes with top 5 SHAP values are considered as the key genes. The core modality on which one gene depends indicates that the SHAP value of that gene ranks higher on this modality than on the others.

#### **Hub module of the updated pathway crosstalk network**

The calculation of the sub-network score can be divided into four steps: average pathway crosstalk network matrix calculation, network pruning, sub-network boundary determination, and score calculation. First, according to the trained model, the updated pathway crosstalk network corresponding to each sample was given. For average pathway crosstalk network matrix, we integrated all updated pathway crosstalk network matrices into a matrix by element-wise average and normalization, calculated as follows:

$$\mathbf{P}_{up} = \min - \max(\frac{1}{N} \sum_{n=1}^N \mathbf{P}'_n)$$

, where $n = 1, 2, \dots, N$ is the $n$th sample, $\mathbf{P}'_n$ is the updated pathway crosstalk network matrix of the $n$th sample, min-max is the min-max normalization function.

Then, we performed network pruning, that is, removed larger pathway nodes which contains more than 100 genes in the network. This was done to control the size of the sub-network and avoid evaluation bias caused by excessive emphasis on larger pathways. After network pruning, the average pathway crosstalk network matrix is denoted as $\mathbf{P}'_{up}$.

Next, we defined the sub-network corresponding to each node and its boundary. When the corresponding element of average pathway crosstalk network matrix exceeds a boundary threshold, we defined that there is a link between two pathways, while otherwise there is no link. The boundary threshold is defined according to the data distribution, which is 99.7% quantile of $\mathbf{P}'_{up}$ in TCGA datasets and 99.9% quantile of $\mathbf{P}'_{up}$ in liquid biopsy datasets. Sub-networks are defined as follows:

$$pathlist_1 = [p_{11}, p_{12}, \dots, p_{1j}, \dots, p_{1r}], P'_{up}(p_{give}, p_{1j}) > cut\ off$$

$$pathlist_2 = [p_{21}, p_{22}, \dots, p_{2j}, \dots, p_{2r}], \sum_{p_{1j} \in pathlist_1} sum(P'_{up}(p_{2j}, p_{1j}) > cut\ off) > 0$$

$$SP_{give}(p_{2i}, p_{2j}) = \begin{cases} P'_{up}(p_{2i}, p_{2j}), & \text{if } P'_{up}(p_{2i}, p_{2j}) > cut\ off \\ 0, & \text{if } P'_{up}(p_{2i}, p_{2j}) \leq cut\ off \end{cases}, p_{2i} \text{ and } p_{2j} \in pathlist_2$$

, where $p_{give}$ is the given pathway as central node, $cut\ off$ is the boundary threshold, $pathlist_1$ is the list of pathways neighboring the central node, $pathlist_2$ is the list of pathways of sub-network, and $SP_{give}$ is the adjacency matrix of sub-network.

Then, we calculated sub-network score as the average of SHAP values of all pathways in the sub-network, which is formulated as:

$$score_{sub} = \frac{1}{len(pathlist_2)} \sum_{p_{2j} \in pathlist_2} SHAP_{p_{2j}}$$

Finally, we defined the sub-network with the highest score as the hub module of the updated pathway crosstalk network.

### Supplementary Note 5: Data collection and preprocessing

#### TCGA data

For benchmark testing, we collected TCGA datasets to evaluate classification performance of Pathformer and existing comparison methods in two classification tasks, including cancer early- and late- stage classification, and cancer low- and high- survival risk classification (**Supplementary Fig. 2**).

Firstly, we used the “TCGAbiolinks” package of *R* software to download RNA expression, DNA methylation, DNA CNV and clinical data of TCGA datasets. Among these downloaded datasets, the RNA expression values were read counts processed by *STAR* and normalized by TPM, the CpG site levels of DNA methylation data are β-values measured using the Infinium HumanMethylation450 BeadChip, and the DNA CNV data were masked copy number segment and gene level score processed by *Gistic2*.

Next, we added label information for classification experiments. For cancer early- and late- stage classification, we defined stage I and stage II as the early stage and stage III as the late stage according to the "pathologic stage" information in clinical data. For cancer low- and high- survival risk classification, we defined samples from patients with survival time greater than 1825 days as low-risk samples and those less than 1825 days as high-risk samples.

Finally, we conducted additional filtering, retaining only those samples that included RNA expression, DNA methylation, DNA CNV, and their corresponding clinical labels in each cancer dataset. We specifically selected cancer datasets with a substantial number of samples, including those for stage classification with over 300 samples and those for survival classification with over 150 samples. Specifically, cancer early- and late- stage classification task involves 10 datasets from TCGA term, including breast cancer (BRCA), neck cancer (HNSC), low-grade gliomas (LGG), bladder cancer (BLCA), melanoma (SKCM), kidney clear cell carcinoma (KIRC), lung adenocarcinoma (LUAD), lung squamous cell carcinoma (LUSC), liver cancer (LIHC) and pan-cancer. The pan-cancer dataset contains 5610 samples, covering 21 cancer types (BRCA, COAD, HNSC, KICH, KIRC, KIRP, LUAD, LUSC, READ, STAD, ACC, BLCA, CHOL, ESCA, LIHC, MESO, PAAD, SKCM, TGCT, THCA, UVM, abbreviation of cancer type according to the TCGA terms). Cancer low- and high- survival risk classification task involves 10 datasets from TCGA, including BRCA, BLCA, SKCM, stomach cancer (STAD), KIRC, LUAD, LUSC, LIHC, thyroid cancer (THCA) and pan-cancer. The pan-cancer dataset contains 3447 samples, covering 33 cancer types (BRCA KIRC, HNSC, LGG, LUAD, LUSC, STAD, BLCA, LIHC, SKCM, THCA, KICH, COAD, KIRP, READ, UCEC, ACC, CESC, CHOL, DLBC, ESCA, GBM, LAML, MESO, OV, PAAD, PCPG, PRAD, SARC, TGCT, THYM, UCS, UVM, abbreviation of cancer type according to the TCGA terms).

##### ***Liquid biopsy data***

To further verify the effectiveness of Pathformer in cancer diagnosis, we collected two complex body fluid datasets from different blood components: the plasma dataset (comprising 98 healthy donors, 90 colorectal cancer (CRC), 23 esophageal cancer (ESCA), 71 STAD, 57 LIHC, and 34 LUAD assayed by total cell-free RNA-seq<sup>3,4</sup> and the platelet dataset (comprising 286 healthy donors, 462 LUAD, 42 CRC, 40 GBM, 39 BRCA, 35 PAAD, and 14 LIHC from two studies assayed by tumor-educated blood platelet RNA-seq<sup>5,6</sup>). For body fluid datasets, we used seven modalities at the RNA level as Pathformer's input, including RNA expression, RNA splicing, RNA editing, RNA alternative promoter (RNA alt. promoter), RNA allele-specific expression (RNA ASE), RNA single nucleotide variations (RNA SNV), and chimeric RNA.

We used a bioinformatics pipeline to preprocess raw sequence reads into datasets of different modalities. Firstly, we used *cutadapt* tool to trim adaptors and low-quality reads, and then removed the reads which can be mapped to ERCC's spike-in sequences, NCBI's UniVec sequences (vector contamination), and human rRNA sequences by *STAR* software. Next, we applied *STAR* software to map all the retained unmapped reads to the hg38<sup>7</sup> genome index built with the GENCODE v27<sup>8</sup> annotation and calculated seven modalities at the RNA level based on the mapping result. The details of the calculation processes are as follows: (1) RNA expression data were read counts aggregated to gene by *featureCounts* and were normalized by TPM. (2) RNA alternative promoter data represented transcript isoform abundances quantified by salmon and were normalized by TPM. We only selected isoforms with transcription start sites within 10 bp (sharing the same promoter) and TPMs greater than 1<sup>9</sup>. (3) RNA splicing data were a series of alternative splicing events with the percent spliced-in (PSI) score calculated using *rMATS-turbo*. (4)

As for RNA editing data, editing sites were identified by *GATK ASEReadCounter* based on REDportal<sup>10</sup> and editing ratios of editing sites were defined as allele count divided by total count. (5) In RNA allele-specific expression data, allele-specific expression gene site were identified by *GATK ASEReadCounter* based on SNP sites and allelic expressions (AE, $AE = |0.5 - \text{Reference ratio}|$, $\text{Reference ratio} = \text{Reference reads} / \text{Total reads}$) were calculated for all sites with $\geq 16$ reads<sup>11</sup>. (6) In RNA single nucleotide variations data, *GATK SplitNCigarReads* was used to split intron-spanning reads for confident SNP calling at RNA level. *GATK HaplotypeCaller* and *GATK VariantFilteration* were used to identify and filter alterations. Allele fraction was defined as allele count divided by total count (reference count and allele count). (7) Chimeric RNA data were identified by remapping unaligned reads to chimeric junctions by *STAR-fusion*. Chimera references were based on GTex<sup>12</sup> and ChimerDB-v3<sup>13</sup>.

### **Supplementary Note 6: Details of model training and test**

In this study, we implemented Pathformer's network architecture using the "PyTorch" package in Python v3.6.9 (codes in https://github.com/lulab/Pathformer). For model training and test, we used 2 times 5-fold cross-validation. We implemented model training, hyperparameter optimization and model early stopping on the training set (80%) and tested model on the test set (20%).

When training the model, we applied cross-entropy loss with class-imbalance weight as the label prediction loss, the ADAM optimizer to train Pathformer, and the cosine annealing learning rate method to optimized learning rate. "ADAM optimizer" is implemented by "Adam" function in the "PyTorch" package. The cosine annealing learning rate method is implemented by "CosineAnnealingWarmupRestarts" function in the "PyTorch" package, with the first cycle step size as 15, the cycle step magnification as 2, the number of warmup steps as 5, the decrease rate of learning rate by cycle as 0.9, the minimum of learning rate as $1e-8$, and the maximum of learning rate as optimal value of hyperparameter optimization.

For hyperparameter optimization, we used grid search with 5-fold cross-validation in the training set with the macro-averaged F1 score as the selection criterion. During grid search process, model was trained for 30 epochs. The key hyperparameters of Pathformer are maximum of learning rate ($lr\_max$) $\in [1e-4, 1e-5]$, dropout probability of classification ($c$) $\in [0.3, 0.5]$, and constant coefficient for row-attention ($\beta$) $\in [0.1, 1]$, a total of 8 possible combinations. **Supplementary Fig. 3** shows examples of grid search on breast cancer dataset. **Supplementary** **Table 2** lists results of optimal hyperparameter combination for each dataset. In addition, we implemented hyperparameter optimization process of each TCGA dataset for benchmark testing, and directly set hyperparameters ($lr\_max=1e-5$, $c=0.3$, $\beta=1$) of liquid biopsy datasets for application.

To validate the convergence of Pathformer, we depicted the training loss related to the number of epochs (**Supplementary Fig. 4**). In both TCGA datasets and liquid biopsy datasets with different sample sizes, the total loss rapidly decreased with iteration and eventually converged to a stable state. To prevent overfitting, we employed an early stopping strategy to determine epoch numbers. The early stop strategy refers to stopping training when the macro-averaged F1 score of the validation set (20% of the training set) consecutively decreased more than 1e-2 on 10 epochs. Then we took the model on the epoch before decline as the final model. Additionally, based on the convergence analysis on the liquid biopsy datasets, we set the training epochs to be greater than 100.

### **Supplementary Note 7: Comparison methods**

For benchmarking, we compared three types of multi-modal integration methods: early and late integration methods based on base classifiers, supervised methods in mixOmics, and deep learning-based integration methods. For deep learning-based integration methods, we benchmarked eight representative models, i.e., eiNN, liNN, eiCNN, liCNN, MOGONet, MOGAT, P-NET and PathCNN.

#### ***Early integration methods based on base classifiers***

Early integration methods based on base classifiers refer to methods that splice different modal data and perform classification by support vector machine (SVM), logistic regression (LR), random forest (RF), or gradient boosting tree (XGBoost). Specifically, we first uniformly transform the different modalities to the gene level modal features by a conversion function. For each modal feature, we selected the top 1000 genes with $FDR \leq 0.05$ of the ANOVA as marker genes, which is implemented by “scikit-learn” package of *Python* v3.6.9. When there were less than 200 genes with $FDR \leq 0.05$ of ANOVA, we used P value instead of FDR. When there were less than 20 genes with P value $\leq 0.05$ of ANOVA, we only selected genes by their rank. For $j$th modal feature, the number of marker genes is $N_j$. Then, we concatenated filtered modal features to obtain features with $N_{gs} = \sum_{j=1}^{D_g} N_j$ dimensions as the input of the base classifier. Next, to be consistent with Pathformer, we divided the dataset into the training set (80%) and the test set (20%) hierarchically, and performed 5-fold cross-validation on the training set for hyperparameter optimization. SVM was implemented by “SVC” function with kernel='rbf' and $C \in [0.01, 0.1, 1, 10, 100]$ in “scikit-learn” package. LR was implemented by “LogisticRegression” function with solver='liblinear', penalty='l2' and $C \in [0.01, 0.1, 1, 10, 100]$ in “scikit-learn” package. RF was implemented by “RandomForestClassifier” function with max\_depth$\in [10, 50, 100, 200, 500]$ and n\_estimators$\in [10, 50, 100, 200, 500]$ in “scikit-learn” package. XGBoost was implemented by “XGBClassifier” function with learning\_rate=0.5, min\_child\_weight=3, gamma=3, subsample=0.7, scale\_pos\_weight=1, objective$\in$['binary: logistic', 'multi: softprob'], max\_depth$\in [10, 50, 100, 200,$

500] and $n\_estimators \in [50, 100, 200, 500]$ in “XGBoost” package.

#### ***Late integration methods based on base classifiers***

Late integration methods based on base classifiers refer to using SVM, LR, RF, and XGBoost to calculate the classification probabilities of different modality data and take the average probability for prediction. Specifically, we performed preprocessing and marker gene selection on each modality data as described in early integration methods based on base classifiers. We took modal features corresponding to each modality as the input of the base classifier, that is, we established $m$ classifiers corresponding to encoded features of $m$ modalities. Finally, we took the average classification probabilities of $m$ classifiers as the predicted probability. Here, data partitioning, hyperparameter optimization and the implementation of base classifiers are consistent with early integration methods based on base classifiers.

#### ***Supervised methods in mixOmics***

The supervised methods in mixOmics refer to partial least squares-discriminant analysis (PLSDA) and sparse partial least squares-discriminant analysis (sPLSDA). PLSDA uses discriminant analysis to project data into latent structures, aiming to find common information across multi-modal data and differentiate between different phenotype groups. sPLSDA is PLSDA appended with sparse regularization. We implemented the PLSDA module and the sPLSDA by the “mixOmics” package of *R* software. Specifically, we performed preprocessing and marker gene selection on each modality data as described in ‘Early integration methods based on base classifiers’ section. Then gene embedding corresponding to each modality were used as input of PLSDA and sPLSDA for model training and testing. In addition, to be consistent with other models, we also divided the dataset into the training set (80%) and the test set (20%) hierarchically, and performed 5-fold cross-validation on the training set to optimize the number of components. The value range of the number of components is [2, 5, 10].

#### ***eiNN and liNN***

eiNN and liNN are early and late integration methods based on fully connected neural network (FCNN). FCNN usually consists of an input layer, multiple hidden layers, and an output layer. eiNN means flattening all modal features (each dimension of gene embedding) and concatenating them into a vector as input for the neural network. liNN means taking modal features corresponding to each modality as separate inputs to the sub neural network, then connecting the output layers together, followed by a fully connected layer for the final output. We used the code on DL-mo’s Github library (https://github.com/zhenglinyin/DL-mo) to implement models. For eiNN, we set 3 hidden layers with dimensions of 500, 100, and 50, dropout rate as 0.1, and lr as $1e-5$. For liNN, we set 3 subnetworks with 2 hidden layers of 100 and 50 dimensions, classification layer with 3 hidden layers dimensions of 100, 50, and 10,

dropout rate as 0.1, and lr as 1e-5. Specifically, we performed preprocessing and marker gene selection on each modality data as described in early integration methods based on base classifiers. For model training and test, we divided each dataset into the training set (80%) and the test set (20%) hierarchically, and used 20% of the training set as the validation set to achieve early stopping of the model. When the number of epochs was more than 200 and the macro-averaged F1 score of the validation set consecutively decreased more than 1e-2 on 10 epochs, we stopped training and took the model on the epoch before decline as the final model.

#### *eiCNN and liCNN*

eiCNN and liCNN are early and late integration methods based on convolutional neural network (CNN). eiCNN means flattening all modal features and concatenating them into a vector as input for CNN. liCNN means taking modal features corresponding to each modality as separate inputs to each CNN, then connecting the output layers together, followed by a fully connected layer for the final output. We used the code on DL-mo's Github library (https://github.com/zhenglinyi/DL-mo) to implement models. For eiCNN, we set 2 CNN layers with kernel size of 1000 and 50, 2 maximum pooling layers with size of 100 and 10, fully connected layer with dimensions of 50, and lr as 1e-5. For liCNN, we set 3 subnetworks with a CNN layer and a maximum pooling layer, classification layer with 3 hidden layers dimensions of 100, 50, and 10, dropout rate as 0.1, and lr as 1e-5. In addition, we performed preprocessing, marker gene selection on each modality data, data partitioning, and early stopping strategy of eiCNN and liCNN are consistent with eiNN and liNN as described above.

#### *MOGONet and MOGAT*

MOGONet model first uses graph convolution to learn weighted sample similarity network and the matrix of a single modality, and then uses the View Correlation Discovery Network (VCDN) to integrate the classification probabilities of different modalities. MOGAT model builds upon MOGONet by replacing the graph convolutional neural network with a graph attention neural network. We used the code on MOGONet's Github library (https://github.com/txWang/MOGONET) to implement MOGONet model and DL-mo's Github library (https://github.com/zhenglinyi/DL-mo) to implement MOGAT model. We set num\_epoch\_pretrain as 200, adj\_parameter as 2, dim\_he as 100, lr\_e\_pretrain as 1e-3, lr\_e as 5e-4, lr\_c as 1e-3, and lr as 1e-5 respectively. We then took modal features corresponding to each modality as the input of MOGONet for downstream analysis. In addition, we performed preprocessing, marker gene selection on each modality data, data partitioning, and early stopping strategy of MOGONet and MOGAT are consistent with eiNN and liNN as described above.

#### *P-NET*

P-NET model is a sparse neural network integrating multiple molecular features based on a multilevel view of biological pathways. In P-NET, all dimensions of gene multi-modal embedding are connected together as inputs, and then distributed on node layers representing a set of genes using weighted links. The other hidden layers of P-NET are constructed based on the hierarchical structure of pathways. The connections between different layers are limited to the child-parent relationship between features, genes, and pathways. In particular, P-NET does not need feature selection. We rewrote the code based on the PyTorch following the code in the P-NET's Github library (https://github.com/marakeby/pnet_prostate_paper) to implement the model, and set pathway dataset to Reactome dataset, n\_hidden\_layers as 5, activation as 'tanh', kernel\_initializer as 'glorot\_uniform', bias\_initializer as 'zeros', batch\_normal as 'False', repeated\_outcomes as 'True', dropout as [0.5, 0.1, 0.1, 0.1, 0.1, 0.1, 0.1], and lr as 1e-5. In addition, data partitioning and early stopping strategy of P-NET are consistent with eiNN as described above.

##### ***PathCNN***

PathCNN model is a classic model that introduces known pathway knowledge for multi-modal integration. PathCNN first uses principal component analysis (PCA) to integrate multi-modal data into the pathway level as pathway images of multi-modal, and then uses convolutional neural network (CNN) to extract high-dimensional features for downstream classification tasks. We used the code on PathCNN's Github library (https://github.com/mkskpi/PathCNN) to implement the model, and set lr to 1e-5. Specifically, we first uniformly transform the different modalities to the gene level modal features by a conversion function. Then, we filtered the modal features corresponding to genes of 146 pathways in PathCNN, and used the PCA to integrate these to obtain multi-modal pathway images. Finally, we took pathway image of each modality as input to PathCNN for downstream analysis. In addition, data partitioning and early stopping strategy of PathCNN are consistent with eiNN and liNN as described above.

##### **Supplementary Note 8: Feature selection for cancer noninvasive diagnosis**

To understand the necessity of liquid biopsy multi-modal integration, we first calculated seven RNA-level modalities as Pathformer's input separately. From results of 2times 5-fold cross-validation in **Supplementary Fig. 8**, we found that the model with all modalities as input had the best comprehensive performance on two datasets, followed by RNA expression-only model and RNA alt. promoter-only model, and some models with other modalities exhibited great fluctuations on different datasets. In order to effectively integrate information without redundancy, we performed further feature selection based on different modality combinations evaluated by Pathformer. First, we calculated the contributions of each modality and its corresponding statistical indicators (**Supplementary Fig. 9a**). Similar to the results of cross-validation, RNA expression was the core modality across

all datasets. Next, we performed 5-fold cross-validation find an optimal modality combination for each dataset (**Supplementary Fig. 9b**). We found that plasma dataset with 7 modalities and platelet dataset with 3 modalities (RNA expression, RNA alternative promoter, and RNA splicing) obtained the best performance.
